# Label-free composition determination for biomolecular condensates with an arbitrarily large number of components

**DOI:** 10.1101/2020.10.25.352823

**Authors:** Patrick M. McCall, Kyoohyun Kim, Martine Ruer-Gruß, Jan Peychl, Jochen Guck, Anthony A. Hyman, Jan Brugués

## Abstract

Biomolecular condensates are membrane-less organelles made of multiple components, often including several distinct proteins and nucleic acids. However, current tools to measure condensate composition are limited and cannot capture this complexity quantitatively, as they either require fluorescent labels, which we show can perturb composition, or can distinguish only 1-2 components. Here, we describe a label-free method based on quantitative phase microscopy to measure the composition of condensates with an arbitrarily large number of components. We first validate the method empirically in binary mixtures, revealing sequence-encoded density variation and complex aging dynamics for condensates composed of full-length proteins. In simplified multi-component protein/RNA condensates, we uncover a regime of constant condensate density and a large range of protein:RNA stoichiometry when varying average composition. The unexpected decoupling of density and composition highlights the need to determine molecular stoichiometry in multi-component condensates. We foresee this approach enabling the study of compositional regulation of condensate properties and function.

## INTRODUCTION

Many compartments in living cells exist as condensed phases of biopolymers, termed biomolecular condensates, which are demixed from the surrounding cytoplasm or nucleoplasm ^1,2^, and are implicated in a wide range of cellular processes ^3^. Phase separation of a simple binary mixture of a polymer in solvent results in a dilute phase coexisting with a polymer-rich condensed phase (**Fig. 1a**). Although demixing of a single full-length protein in a binary mixture is often sufficient to reconstitute simplified condensates ^4–7^, condensates *in vivo* contain dozens of components ^8–10^. Indeed, the functional identity of a particular condensate inside a cell is determined by its composition. Unlike binary systems, such multi-component condensates possess a continuum of compositions ^11^, each connected to a coexisting dilute phase via a tie-line in a phase diagram (**Fig. 1b**). Changes in component abundance can thus shift the system to a new tie-line, altering condensate composition and physical properties ^12,13^. Despite its central role in physically defining condensates and specifying their properties, the composition of multi-component condensates *in vivo*, and component stoichiometries in reconstituted systems *in vitro*, are largely unknown.

**Fig 1:**
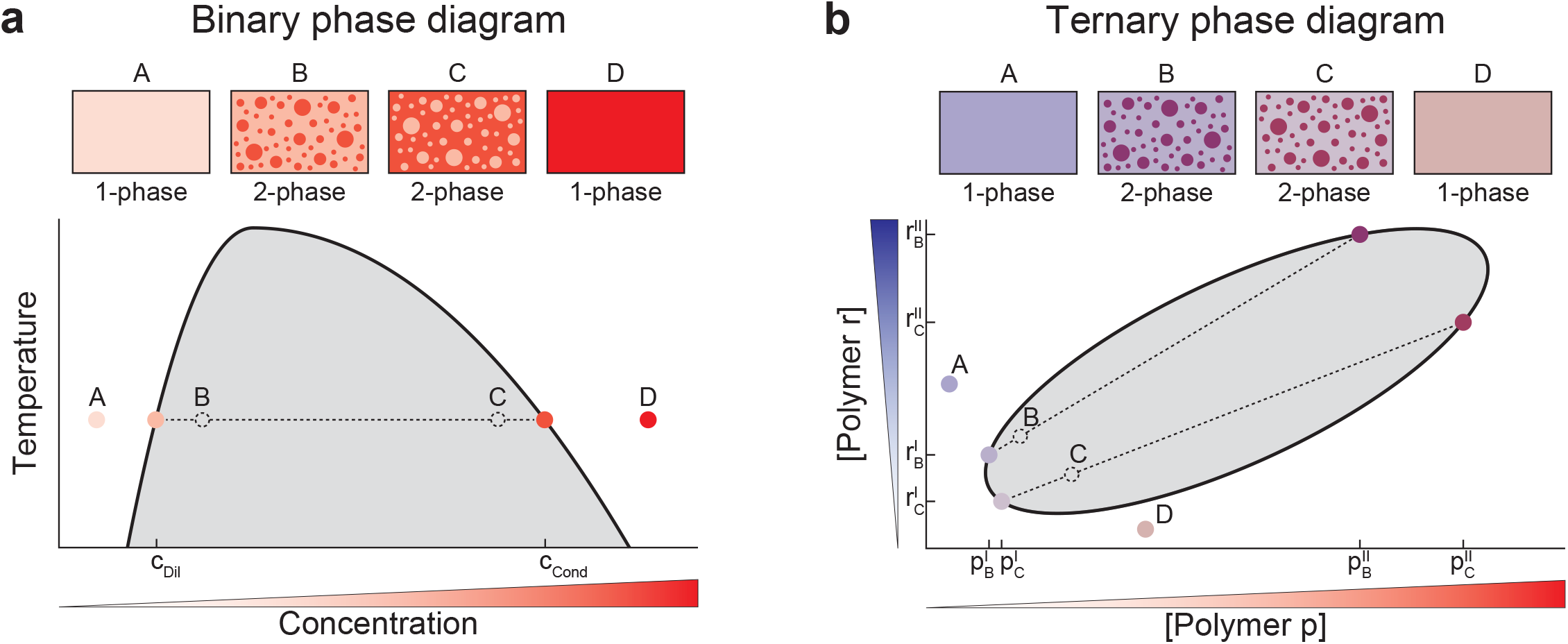
Phase diagrams of binary and ternary mixtures. **a**, Phase diagram in a prototypical binary mixture with sketches of samples prepared in the 1-phase (A,D) and 2-phase regimes (B,C). Compositions of coexisting phases, *c*_*Dil*_ and *c*_*Cond*_, lie on the binodal curve (solid black) that separates the 1-phase (mixed) and 2-phase (demixed) regimes. A tie-line (dashed black) connects compositions of coexisting phases to the corresponding average composition (open dashed black circle). Varying average solute concentration in the 2-phase regime (gray) changes the relative volumes of coexisting phases but not their composition. **b**, Phase diagram in a prototypical ternary mixture with sketches of samples prepared at different average compositions. Unlike the binary case, ternary mixtures prepared at different points in the two-phase regime may lie on different tie-lines, yielding compositionally distinct pairs of coexisting phases.

In multi-component systems, condensate composition is typically estimated by fusing each molecular species to a spectrally-distinct fluorescent tag ^12,14,15^. Although powerful, fluorescent tags contribute to interactions between species and with solvent, potentially shifting the thermodynamic balance that specifies phase composition ^16^. Interactions with the condensed phase can also drive strong deviations in fluorophore characteristics relative to behavior calibrated in the dilute phase ^17,18,15^, confounding quantification. Label-free techniques avoid these issues entirely, but existing approaches require harsh treatments and are effectively limited to binary systems for native-like molecules. For example, traditional bulk approaches like ultra-violet (UV) absorption ^20,21^ and thermogravimetric analysis ^22^ require condensate dissolution or destruction and impose sample requirements that are often inaccessible with the modest yields obtained by recombinant expression and purification of endogenous cellular condensate components (**Supplementary Note 1**). Though confocal Raman spectroscopy enables measurements of intact condensates ^18,19^, the experimental requirements of high laser exposure or nanoparticles for surface-enhancements may alter condensate dynamics, and a calibration sufficient to resolve multi-component composition has yet to be demonstrated.

Reflecting these limitations, recent UV absorption measurements of the composition of condensates reconstituted from recombinant proteins looked exclusively at binary systems with intrinsically disordered protein regions (IDRs) rather than the full-length proteins ^23–25^. This is in part because those fragments could be purified with sufficient yield from bacteria following denaturation. That is not an option in many cases, however, as bacteria lack the machinery needed to add post-translational modifications (PTMs) or assist folding of certain protein domains, and denaturants may irreversibly alter protein conformational ensembles. Thus, it remains unclear how the additional features of native-like proteins, including PTMs, native-like conformational ensembles, and the additional domains present in full-length proteins, will change the picture now emerging for condensates formed from IDRs alone. As functional roles for PTMs and structured protein domains accumulate alongside molecular parts lists ^8–10^, there is a pressing need for methods to reveal condensate composition in more faithful reconstitutions that include multiple native-like components.

Here, we present a label-free method to precisely measure the composition of micron-sized condensates reconstituted from an arbitrary number of components. Overall, this method requires 1000-fold less material than bulk label-free alternatives, which enables dynamic and temperature-dependent measurements of condensates formed from full-length native-like components. We demonstrate that quantitative phase imaging (QPI) can be used to extract refractive index differences between demixed phases, ∆*n*, which we convert to condensed-phase concentration for binary systems. By combining these ∆*n* with tie-line measurements, we then show that the concentrations of an arbitrary number of individual species can be resolved quantitatively in multi-component condensates. We demonstrate this explicitly for a model ternary system of full-length FUS protein and RNA, and reveal unexpected features in the composition of these multi-component condensates. Additionally, we find that protein concentrations vary from 87 to more than 470 mg/ml in condensates depending on sequence and use of labels, and uncover a dramatic increase in density with age that may underly previous reports of mechanical hardening ^26^. By resolving the chemical composition of multi-component condensed phases *in situ* and in unprecedented detail, we anticipate that this label-free method will enable mechanistic studies of complex composition regulation of biomolecular condensate properties and function with multiple native components.

## RESULTS

### Optical concept

To measure the compositional difference between micron-sized droplets and the coexisting dilute phase, we employ quantitative phase imaging (QPI) ^27,28^. Physically, QPI measures the optical phase shift accumulated along a wavefront as it traverses spatial inhomogeneities in refractive index within a sample, such as high-refractive index droplets immersed in a lower-index medium **(Fig. 2a)**. Within the first-order Born approximation ^29^, the optical phase shift ∆*φ* measured by QPI at pixel (*x, y*) is proportional to the product of refractive index difference, ∆*n*, and droplet shape

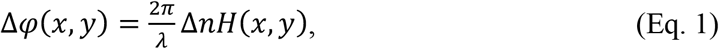

where λ is the imaging wavelength and *H*(*x, y*) is the projected height of the droplet along the imaging axis. Thus, the refractive index difference between coexisting thermodynamic phases can be obtained from the droplet‘s shape and optical phase shift. The refractive index of an aqueous protein solution is, in turn, a linear function of concentration over a wide range ^30,31^. In this regime (see **Supplementary Note 2**), the condensed-phase protein concentration, *c*_*Cond*_, in a binary system is given by

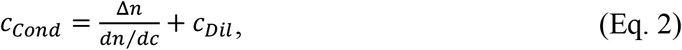

where *c*_*Dil*_ is the dilute phase concentration and *dn/dc* is the slope of the concentration-dependence of refractive index, which can be estimated from amino acid sequence ^32^. This suggests QPI is suitable to measure the concentration difference between a condensate and its coexisting dilute phase in a binary system when condensate shape is known.

**Fig 2:**
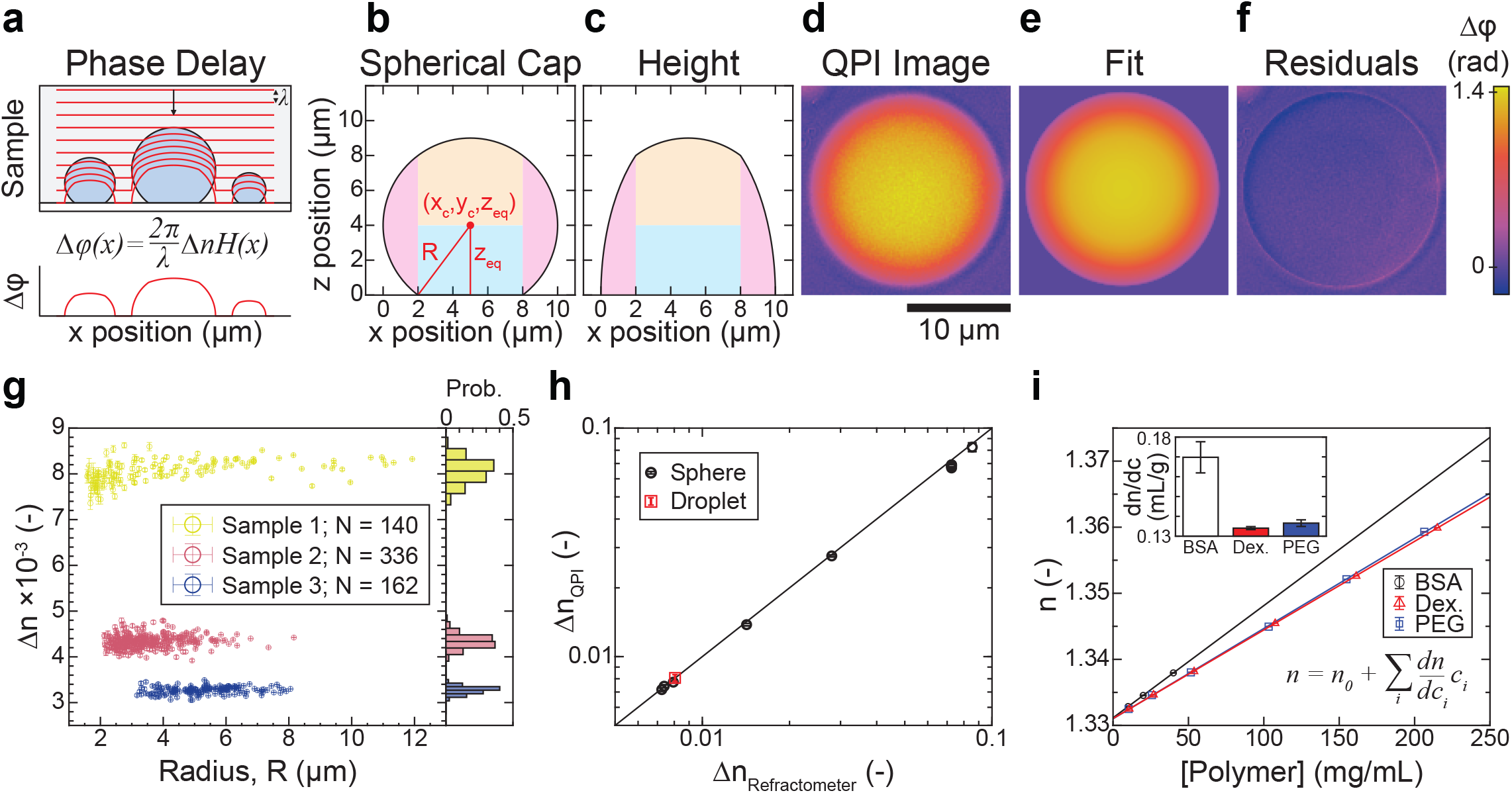
QPI measures the refractive index of micron-sized droplets *in situ*. **a**, Schematic of optical wavefronts (red) distorted by droplets on a flat surface (top) and the cumulative optical phase shift (bottom). **b**, Schematic defining the geometry of a spherical cap. **c**, Projected height profile *H(x)* for the geometry in **b**. Shading in **b**,**c** denotes separate terms in the analytic expression for *H(x)*, (see Methods). **d**, Phase image of dextran-rich droplet on passivated glass. **e**, Fit of droplet in **d** to spherical cap. **f**, Residuals from fit. Scalebar in **e**-**f** is 10 µm. **g**, Refractive index difference ∆n vs. droplet radius *R* extracted from fits to individual dextran-rich droplets obtained from 3 different PEG/Dextran mixtures (colors). Error bars are 95 % confidence intervals. Probability histograms at right. **h**, Refractive index differences measured with QPI vs. bulk refractometry for PEG/Dextran mixtures (red) or silica spheres in different glycerol-water mixtures (black). Error bars are standard deviation. Solid line is *y = x*. **i**, Refractive index is a linear function of component concentration for three model biopolymers (BSA, Dextran, PEG). Inset: Slopes (*dn/dc*) for each polymer with 95 % confidence intervals.

### Measuring droplet shape on flat surfaces

To characterize the shape of condensates typical of *in vitro* reconstitution experiments, we model them physically as sessile fluid droplets on a flat substrate (**Fig. 2a)**. In this context, droplet shape is determined at equilibrium by interfacial tension opposing gravitational settling of the denser fluid ^33^. For droplets smaller than the capillary length *l*_*c*_ = (γ/∆*ρg*) ½, the interfacial tension γ is sufficiently strong to suppress expansion of the interfacial area driven by gravitational effects and the resulting shape is described to an excellent approximation as a spherical cap ^33^ (**Fig. 2b**). This approximation is valid for most reconstituted condensates, which are typically smaller than our capillary length estimate of ≳ 30 µm (**Extended Data Fig. 1**). In this limit, four parameters suffice to fully characterize droplet shape: droplet radius *R*, the position of the droplet center (*x*_0_, *y*_0_), and the height of the equatorial plane above the substrate, *Z*_*eq*_ (**Fig. 2b, Methods**). Importantly, the corresponding projected height function *H*(*x, yi|R, x_0_, y_0_, Z_eq_*) has a closed analytic form that we use to determine the refractive index and shape parameters of droplets from QPI images by fitting (**Fig. 2c, Methods**).

### Experimental validation and conversion from ∆*n* to concentration

To validate the approach experimentally, we use two different reference systems. The first is a well-characterized PEG/Dextran aqueous two-phase system ^34^ for which we can readily measure ∆*n* independently by bulk refractometry (**Methods**). QPI images of Dextran-rich droplets on passivated coverglass and surrounded by a coexisting PEG-rich phase (**Fig. 2d**) are well-modeled as spherical caps (**Fig. 2e**), as evidenced by small and spatially unstructured residuals in the droplet interior following fitting (**Fig. 2f**). For the best fits, ∆*n* is independent of size for three different PEG/Dextran compositions (**Fig. 2g, left)**, and approximately symmetrically distributed (**Fig. 2g, right**), indicative of equilibrated phases and uncertainty dominated by statistics rather than systematics, respectively. Crucially, the ∆*n* values extracted from fL-droplets in QPI images are in excellent agreement with those measured independently from 100-µL volumes of each phase using a digital refractometer (**Fig. 2h**). To validate application of the first-order Born approximation for larger ∆*n*, we used silica microspheres suspended in glycerol-water mixtures as a second reference system, where ∆*n* is set by the glycerol/water ratio. As with the dextran droplets, we recovered the expected shape without bias and find excellent agreement between ∆*n* measurements extracted from QPI images and those expected on the basis of digital refractometry measurements, now over a much larger range (**Fig. 2h, Extended Data Fig. 2**). Taken together, these data demonstrate that the present analysis pipeline extracts accurate geometric and optical measurements of homogeneous sessile droplets from QPI image data for ∆*n* of at least 0.085.

Conversion from ∆*n* to compositional differences requires knowledge of the refractive index increments, *dn/dc*, for each partitioning component. Using bulk refractometry, we determined *dn/dc* from the concentration-dependence of solution refractive index for several representative (bio)polymers (**Fig. 2i, Extended Data Fig. 3**). In each case, we found excellent linearity over the entire range probed. Importantly, the measured dn/dc value for BSA is consistent with estimates from amino acid sequence ^32^. This validates the use of sequence-based dn/dc estimates in the following, particularly given the impracticality of direct measurement for many recombinant proteins (**Supplementary Note 1**). With *dn/dc* estimated from protein sequence, ∆*n* measured by QPI, and *c*_*Dil*_ measured by standard analytical methods or neglected (**Supplementary Note 3**), Eq. 2 enables calculation of condensed-phase protein concentrations in binary systems.

### Condensates of native-like proteins

To demonstrate the suitability of our method for condensates reconstituted with native full-length proteins, we first investigated PGL-3, which is a major component of P granules in *C. elegans* ^8^. PGL-3 forms condensates *in vitro* under low salt ^6^. Using QPI, (**Fig. 3a**), we found the concentration in individual PGL3 condensates is symmetrically distributed about a mean of 87.0 ± 0.1 mg/ml (s.e.m., N = 269) (**Fig. 3c**), approximately 1000-fold higher than that in the coexisting dilute phase. The standard deviation of the measured population is only 1.7 mg/ml (**Fig. 3c**), yielding a low coefficient of variation (1.9 %) which reflects the high precision of the QPI method as well as the low droplet-to-droplet variation expected near phase equilibrium. Given the impracticality of bulk label-free measurements for this untagged full-length protein (**Supplementary Note 1**), we employed Optical Diffraction Tomography (ODT) for comparison (**Fig. 3b,c**). ODT is a related approach recently applied to stress granules ^10,35^. We find ODT to provide a comparable accuracy to QPI, though with reduced precision (**Fig. 3c**).

**Fig 3:**
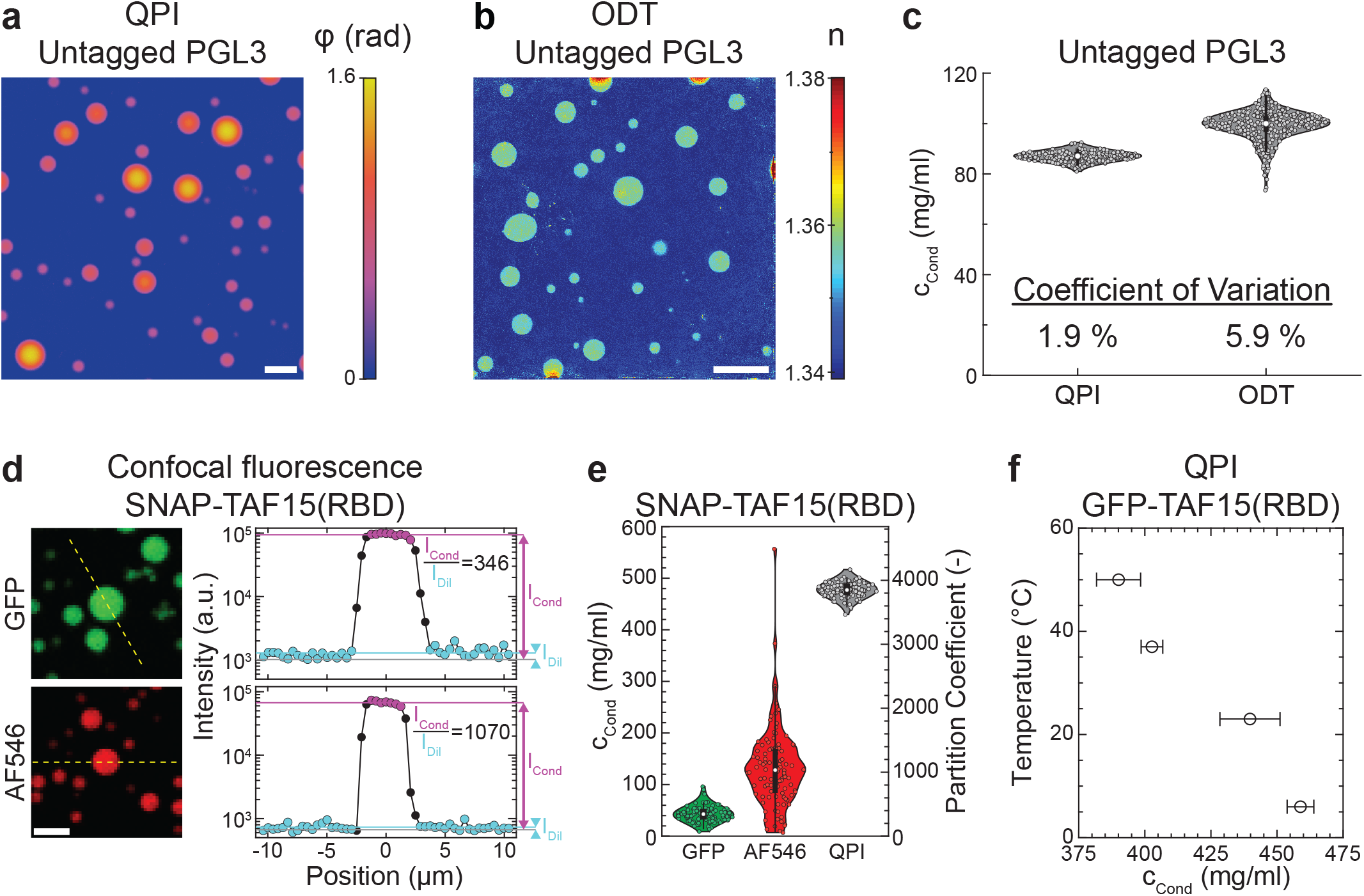
Label-free composition measurement of multi-domain protein condensates. Quantitative phase, **a**, and refractive index, **b**, images of untagged full-length PGL3 condensates acquired by QPI and ODT, respectively. Scalebar is 10 µm. **c**, PGL3 concentration measured from individual condensates by QPI (N = 269) or ODT (N = 355). **d**, Confocal fluorescence images of SNAP-TAF15(RBD) condensates doped with either 23 % mEGFP-TAF15(RBD) or 12 % AlexaFluor546-SNAP-TAF15(RBD). Scale bar is 5 µm. Intensity profiles along the dashed yellow lines are shown at right. Gray, cyan, and lavender lines denote the average detector background, dilute-phase intensity, and condensed-phase intensity. **e**, Comparison of SNAP-TAF15(RBD) concentrations in condensates (left) and partition coefficients (right) measured by QPI (N = 119) or confocal fluorescence intensity ratios of mEGFP (N = 107) or AF546 (N = 104). White circles denote medians, thick black bars are the interquartile range, and whiskers extend 1.5x beyond the interquartile range. Protein concentration in the dilute phase is taken to be 1.97 µM. **f**, Temperature-dependence of protein concentration in mEGFP-TAF15(RBD) condensates (i.e. condensed-branch of binodal) measured by QPI. Error bars are s.e.m.

We next formed condensates using constructs derived from full-length FUS and the RNA-binding domain of TAF15, TAF15(RBD), both reported previously in ^7^, and measured their composition with QPI. Interestingly, we find these condensates to be much denser than those of PGL3 (**Table 1**), with the 34% polymer volume fraction in TAF15(RBD) condensates comparable to that in protein crystals ^36^. Taken together, these data not only reveal that protein sequence can tune condensate composition over at least a 5-fold range, but also demonstrate that the QPI method enables precise label-free measurements on condensates of full-length proteins. This removes the primary practical barrier preventing study of native full-length proteins that are available only in limited quantities but which are most physiologically relevant.

**Table 1.** Compositions of biomolecular condensates in vitro.

| Construct | Conditions<br>(T in °C, [salt], pH) | $c_{\text{cond}}$<br>mg/ml<sup>a</sup> | Protein Volume<br>Fraction<sup>b</sup> | Partition<br>Coefficient | Method |
| --- | --- | --- | --- | --- | --- |
| PGL3 | (25.0, 75 mM, 7.4) | $87.0 \pm 1.7$ | $0.0649 \pm 0.0013$ | n.d. | QPI |
| PGL3 | (21.5, 87 mM, 7.4) | $99.2 \pm 5.9$ | $0.0740 \pm 0.0044$ | n.d. | ODT |
| PGL3-mEGFP | (21.5, 87 mM, 7.4) | $113.2 \pm 8.6$ | $0.0844 \pm 0.0064$ | n.d. | ODT |
| SNAP-TAF15(RBD) | (37.0, 100 mM, 7.4) | $477.1 \pm 13.7$ | $0.3400 \pm 0.0098$ | 3850<sup>c</sup> | QPI |
| FUS-mEGFP | (21.0, 150 mM, 7.4) | $337.3 \pm 8.2$ | $0.2395 \pm 0.0058$ | 860<sup>d</sup> | QPI |

<sup>a</sup> Uncertainty represents standard deviation from a population of at least 100 individual condensates
<sup>b</sup> Fraction of the condensed phase volume occupied by protein, $\phi \equiv c_{\text{cond}} \bar{v}$ ; $\bar{v} \approx 0.75$ mL/g for PGL3 constructs and $\bar{v} \approx 0.71$ mL/g for TAF15 and FUS constructs
<sup>c</sup> $c_{\text{Dil}} = 1.97 \pm 0.09$ $\mu\text{M}$ , $M_w = 62.92$ kDa; <sup>7</sup>
<sup>d</sup> $c_{\text{Dil}} = 4.87 \pm 0.48$ $\mu\text{M}$ , $M_w = 80.38$ kDa; <sup>7</sup>

### Influence of fluorescent labels

Conventional approaches to study condensates of full-length proteins typically require fluorescent labels, introducing two complications in phase-separating systems. First, large GFP-or SNAP-tags may alter the same phase behavior they are used to measure by shifting the balance of polymer-solvent interactions that drive demixing. Second, fluorophore photo-physics may vary between the starkly different chemical environments presented by the two phases (**Table 1**). We leveraged our label-free method to assess both of these potential effects. In the case of PGL3, we found that fusion to an mEGFP-tag increases the protein mass concentration in the condensed phase by 14 % (**Table 1**). As the tag increases the construct’s molecular weight by more than 14 %, this actually corresponds to a decrease in the molar protein concentration, consistent with the tag imparting a modest solubilizing effect.

To test for environmentally-sensitive fluorescence, we used scanning confocal microscopy to measure fluorescence in SNAP-TAF15(RBD) condensates doped with either mEGFP-or AlexaFluor546-SNAP-tagged (AF546) constructs (**Fig. 3d, Methods**). The partition coefficient P ≡ c_Cond_ /c_Dil_ of ∼350 obtained from mEGFP fluorescence suggests a condensed-phase concentration of only 43 mg/ml, underestimating the 477 ± 14 mg/ml value we measure with QPI by over 10-fold (**Fig. 3e**). This indicates that the relationship between fluorescence intensity and concentration differs between phases, and we suspect that enhanced quenching from the high protein concentration in the condensed phase is largely responsible for the decreased quantum yield we infer there ^37^. Surprisingly, assessing partitioning by fluorescence of the more solvent-accessible AlexaFluor546-labeled construct underestimates the concentration by 3.6-fold (**Fig. 3e**). The differential sensitivity we see with different fluorophores suggests that brightness may vary for each dye/condensate pair and be challenging to correct for *a priori*. Taken together, these data demonstrate that fluorescent labels compromise condensate composition measurements in two distinct ways, sometimes dramatically, underscoring the importance of label-free approaches like QPI.

### Temperature-dependent phase behavior

Biomolecular condensates are intrinsically temperature-dependent as thermodynamic phases, making temperature an important control parameter subject to evolutionary selection ^38^. To test whether we could detect temperature-induced composition variation with QPI, we analyzed phase images of TAF15(RBD) condensates acquired at temperatures between 5 and 50 ºC set with a custom temperature stage (**Methods**, ^39^). After accounting for the temperature-dependence of optical constants in Eq. 1 (**Extended Data Fig. 4**), we find that the condensed-phase protein concentration decreases significantly from ∼ 460 to ∼ 390 mg/ml with increasing temperature (**Fig. 3f**). This is indicative of an upper-critical solution temperature (compare to **Fig. 1a**), as has been reported for several other ^23,40^, though not all ^38^, RNA-binding proteins. Finally, the dry objective lenses used for QPI enable fast temperature equilibration by avoiding direct coupling of their thermal mass to the temperature stage.

### Complex aging dynamics in binary systems

Motivated by recent work demonstrating that the mechanical properties of many protein condensates undergo an aging process ^41–43,26^, we hypothesized that there may be a corresponding change in composition as condensates age. To this end, we used QPI to measure the composition of individual PGL3 condensates over 20 hours (**Fig. 4**). During this time period, we observed droplets to noticeably shrink (**Fig. 4a, top, Supplementary Video 1**), as previously shown ^26^. While the shrinkage would be apparent by simple brightfield imaging, QPI indicated that the optical phase shift also increased with time, despite the reduction in droplet size (**Fig. 4a, bottom**). By fitting the QPI data as before, we were able to precisely measure the composition (**Fig. 4b**) and volume (**Fig. 4c**) of individual condensates over this timeframe, revealing surprisingly coordinated dynamics. From these concentration and volume data, we calculated the number of proteins in the condensate (**Fig. 4d**) and found that the nearly 2-fold concentration increase was approximately balanced by a volume decrease, such that the total number of protein molecules in the condensate decreased by only 15 %. These observations indicate that the condensate necessarily expelled a significant amount of solvent while aging.

**Fig 4:**
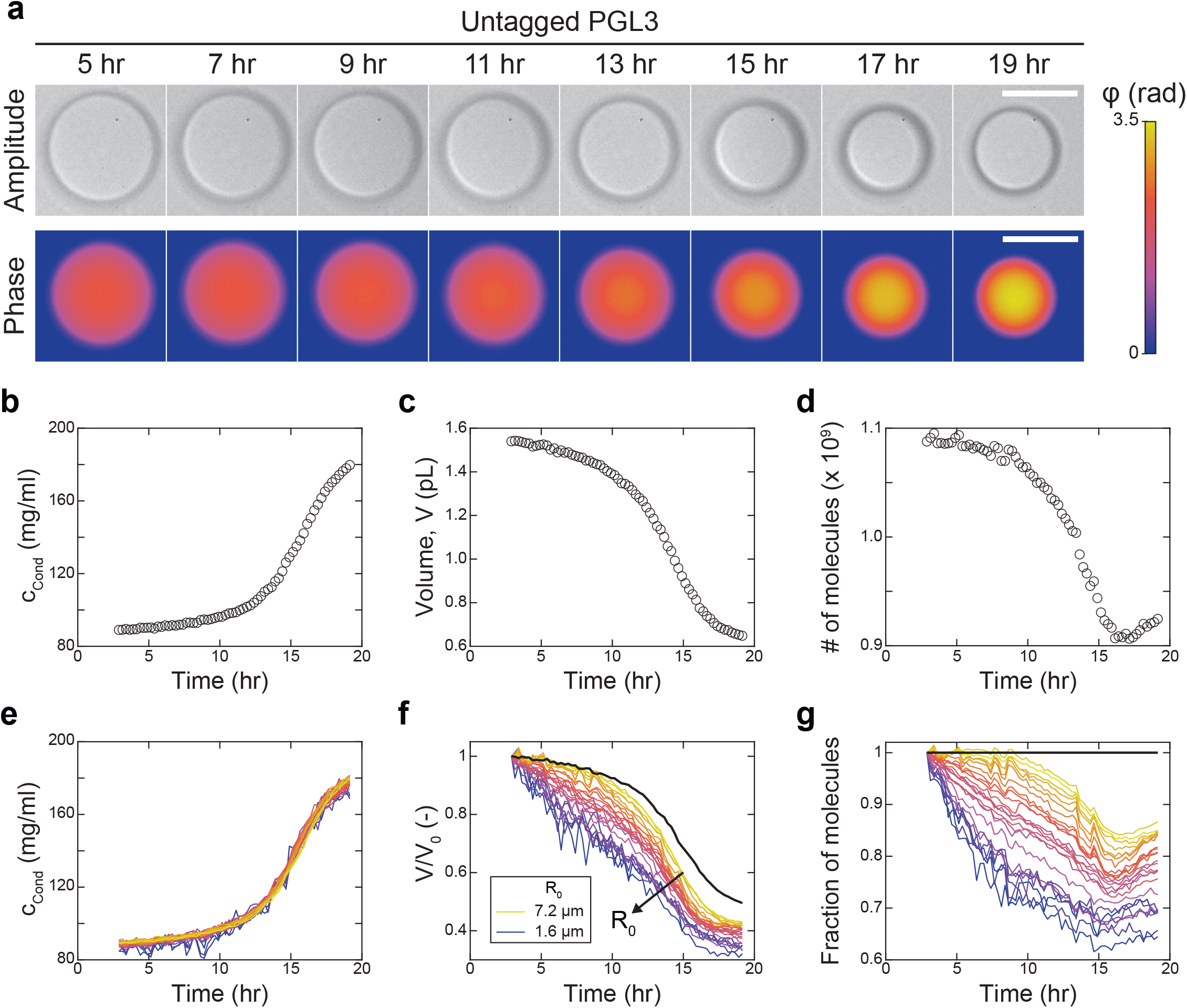
QPI reveals complex aging dynamics in binary systems. **a**, Timelapse of untagged PGL3 condensate (top, QPI amplitude; bottom, QPI phase). Scale bar 10 µm. Concentration **b**, volume **c**, and number of protein molecules in the condensate **d** for the example in **a**. Time is relative to induction of phase separation. **e**, Time-dependence of concentration is identical for N = 25 differently sized condensates. **f**, Normalized volume varies continuously with initial condensate size *R*_*0*_. Initial shrinkage rate decreases with increasing R_0_. **g**, Fraction of molecules in 25 individual condensates over time. Thick black lines in **f**,**g** show volume and molecule count dynamics if protein mass were conserved inside condensates.

To quantify whether this near cancellation was serendipitous for this particular condensate, we analyzed the dynamics of 24 additional condensates with a range of initial sizes over the same period (**Fig. 4e-g**). Strikingly, we found that the kinetics and extent of concentration increase were identical for all condensates, independent of size (**Fig. 4e**). In contrast, the kinetics and extent of the volume decrease both showed systematic size dependencies, with smaller condensates losing volume faster and to greater extent than larger condensates (**Fig. 4f**). As a result, the fraction of molecules retained shows a marked dependence on condensate size, with larger condensates retaining more molecules (**Fig. 4g**). We speculate that the size dependence in the volume kinetics may stem from Ostwald ripening operating in parallel with an additional as yet unknown process driving the contraction and water expulsion. Taken together, these data demonstrate the suitability of QPI to monitor the composition of many individual droplets in parallel, providing insight into the complex interplay between the physical processes driving ripening and aging.

### Measuring binodals and tie-lines with an arbitrary number of components

As condensates in cells are typically enriched in several distinct biomolecular species, we next asked whether QPI could specify the composition of reconstituted condensates with multiple components. In the linear approximation (**Extended Data Fig. 5**), the refractive index difference between two phases with *N* solutes is

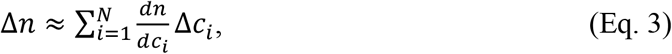

where ∆*c*_*i*_ is the concentration difference of the *i*^th^ component between the two phases. While knowledge of ∆*n* constrains ∆*c* to a single value in the binary systems studied above, the challenge for systems with multiple solutes is that Δ*n* constrains the ∆*c*_*i*_ only to an (*N*-1)-dimensional manifold populated by compositions of equal refractive index. For a ternary system with *N* = 2 solutes and 1 solvent, for instance, this manifold can be visualized as a line in the (*c*_1_, *c*_2_)-plane (**Fig. 5a**). As all compositions along this isorefractive line are compatible with the measured ∆*n*, additional relationships between the ∆*c*_*i*_ are required to uniquely specify condensate composition.

**Fig 5:**
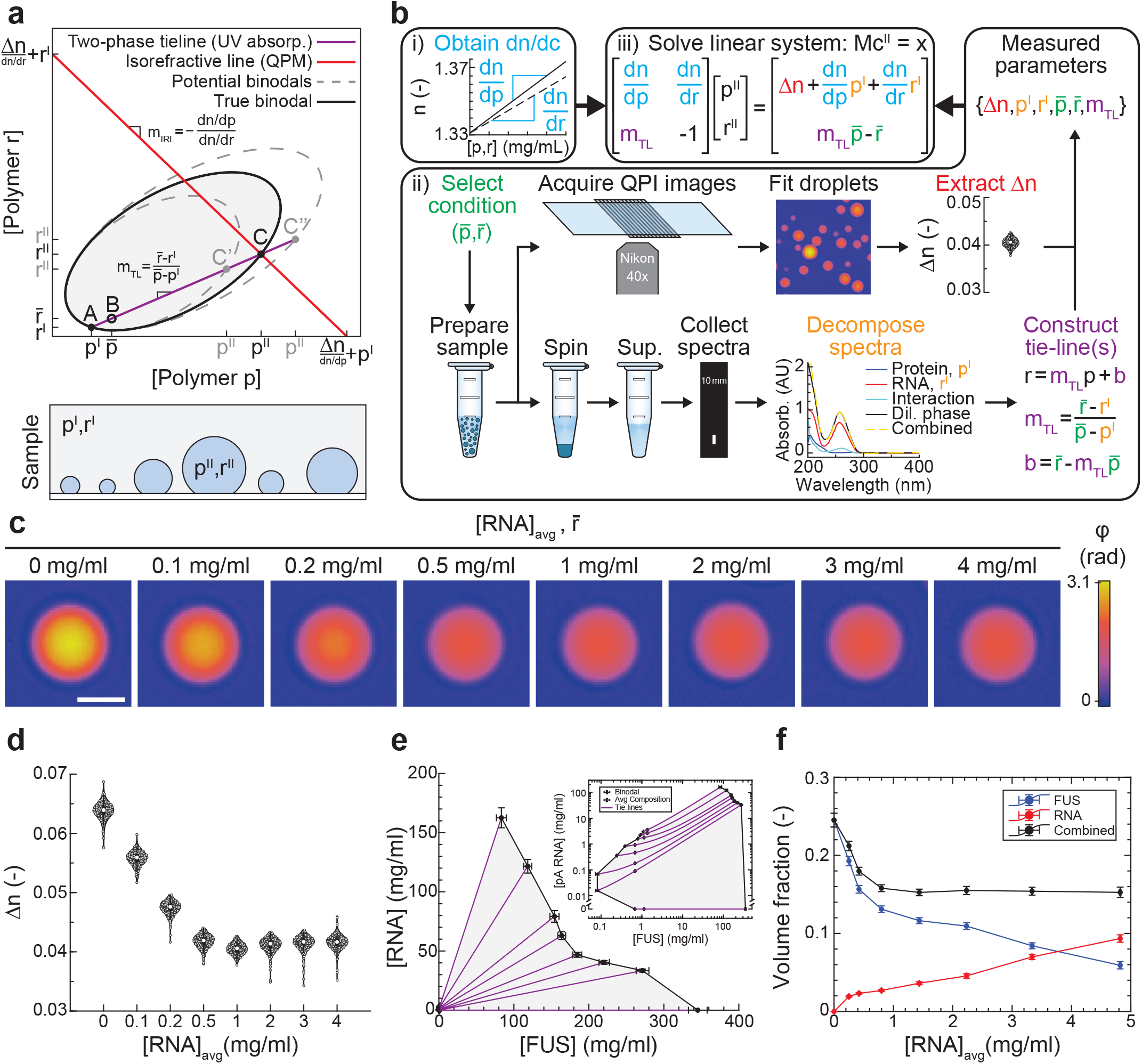
Composition determination of multi-component biomolecular condensates. **a**, Top: Schematic of multi-component measurement approach illustrated for a model ternary system. Condensate composition (*p*^*II*^, *r*^*II*^) is determined by the intersection of the tie-line with the line of constant refractive index (isorefractive line). Bottom: Schematic of coexisting multi-component phases. **b**, Workflow to obtain all parameters in the system of linear equations. i) Refractive index increments are determined once for each partitioning component. ii) For each tested composition 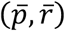 in the two-phase regime, ∆*n* is measured with QPI and the dilute phase composition (*p*^*I*^, *r*^*I*^) is inferred from decomposition of UV-VIS absorbance spectra. The points 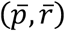 and (*p*^*I*^, *r*^*I*^) suffice to define the tie-line, whose slope *m*_*TL*_ is the final required parameter to solve the linear system, iii). **c**, QPI images of FUS/RNA condensates for a range of RNA concentrations 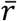 Scalebar is 3 µm. **d**, Measured ∆*n* distributions for the samples in **c. e**, Experimentally-determined ternary phase diagram. Inset shows the same data on loglog axes. Error bars denote standard deviation for dilute-phase and average concentrations, and error propagation via Jacobian for condensed-phase concentrations. **f**, Polymer volume fractions in the condensed phase as a function of 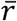 with 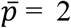 mg/ml. Error bars correspond to the uncertainty in condensed-phase concentration rescaled by polymer partial specific volumes.

Here, we take advantage of the fact that a tie-line connects the total system composition averaged over both phases, 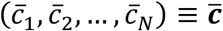 to the compositions of the coexisting phases ***c***^***I***^ and ***c***^*II*^ located on the (*N*-1)-dimensional dilute and condensed binodal manifolds, respectively. For ternary systems, the binodals can be visualized as bounded curves in the (*c*_1_, *c*_2_)-plane (**Fig. 5a**). Mass conservation guarantees that tie-lines are straight (**Fig. 5a, Supplementary Note 4**). Crucially, this provides *N*-1 linearly independent constraints, which can be seen by noting that projections of the tie-line in each of the *N*-1 (*c*_1_, *c*_*i*≠1_)-planes must all be straight. In principle, tie-line constraints of the form 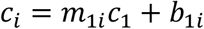 where 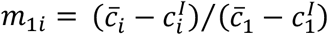 and 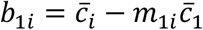 for *i* ≠ 1 can be obtained from knowledge of overall sample conditions, 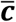, and composition measurements of the abundant dilute-phase 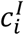 using traditional approaches from analytic chemistry (**Supplementary Notes 3**,**5**). Combined with Eq. 3, these form a linear system of *N* equations of the form *M****c***^*II*^ = ***x***, where the matrix *M* and vector ***x*** contain measured quantities (see **Fig. 5b, Supplementary Note 5** for explicit expressions for ternary and (*N*+1)-component systems, respectively). By solving this system of equations, the composition of the multi-component condensate is written in terms of known optical quantities and concentrations as

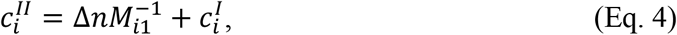

where 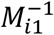 are elements of the system’s matrix inverse (**Supplementary Note 5**). Thus, by leveraging knowledge of tie-lines to provide the missing the constraints, this method is, in principle, capable of resolving the composition of biomolecular condensates with an arbitrary number of components.

### Phase diagram for ternary system with full-length FUS protein and RNA

To validate this approach, we used poly(A) RNA and full length FUS-mEGFP, which localizes to RNA-rich stress granules in eukaryotes ^10^, as solutes in a model ternary system (*N* = 2). Following the workflow in **Fig. 5b**, we prepared systems in the two-phase region for a range of total RNA concentrations. To obtain the dilute-phase composition, we decomposed UV absorbance spectra of the dilute phase into protein and RNA contributions (**Fig. 5b, Methods**). Combined with the system average compositions, the decomposed spectral data produce a set of physically compatible tie-lines (**Extended Data Fig. 6**). To obtain ∆*n*, we analyzed QPI data (**Fig. 5c**), finding that ∆*n* decreases and saturates with increasing total RNA (**Fig. 5d**). Using these data as inputs in Eq. 4 (**Fig. 5b**), we calculated the corresponding points on the condensed binodal branch and plotted these together with the dilute binodal and tie-lines as a full phase diagram (**Fig. 5e**). Reassuringly, this phase diagram captures the re-entrant behavior with increasing RNA inferred previously in related systems without actually measuring the phase boundary ^44,45^. In addition to connecting variation in system average composition directly to its consequences on the condensate, which was seldom accessible previously ^21^, this phase diagram exposes surprising features in the condensed binodal branch, including a kink and linear section (**Fig. 5e**). The ability of this method to resolve the molecular composition of multi-component condensates reveals that rising RNA concentrations are compensated by decreasing protein concentrations such that a constant polymer volume fraction is maintained near cytoplasmic levels (**Fig. 5f**), which would have been challenging to infer using conventional techniques.

## DISCUSSION

Our results show that the present QPI-based method enables quantitative composition measurements of biomolecular condensates reconstituted with full-length proteins in binary and multi-component mixtures. Our finding that a common fluorescent tag can significantly alter condensate composition (**Table 1**) highlights the need for label-free measurement approaches like QPI. As a microscopic method, QPI offers several advantages over traditional label-free strategies based on bulk measurements. The first and most dramatic of these is the ∼1000-fold reduction in sample requirements, which removes the primary barrier to measurements on condensates reconstituted from native full-length proteins (**Fig. 3a,b**). Second, whereas time-resolved measurements of composition in response to changing solution conditions, temperature, or intrinsic sample dynamics are challenging with bulk approaches, QPI’s compatibility with open samples and dry objectives makes it easy to record sample dynamics with second-scale time resolution (**Fig. 4**) and in response to environmental changes (**Fig. 3f**). Third, measuring composition of micron-sized condensates *in situ* provides access to information like size-dependent composition (**Fig. 2g**), homogeneity (**Fig. 2f**), and dynamics (**Fig. 4e-g**) that are not available with bulk techniques. Finally, we note that the high precision demonstrated here with QPI not only enables the method to resolve subtle compositional differences (**Table 1**) and unexpected binodal features (**Fig. 5f**) with high confidence, but could also serve to constrain competing thermodynamic models of condensate formation ^22,7,46,24,47,25^.

An important benefit of the present method is its ability to disentangle the composition of multi-component biomolecular condensates. All endogenous condensates are expected to contain multiple components, with some of the best-characterized known to house dozens ^8–10^. Knowledge of component identity and quantity will be essential for design and interpretation of increasingly faithful reconstitution experiments as well as studies of chemical reactions in synthetic condensates that account for reacting species. Here, we validated our methodology by measuring the full phase diagram for a ternary mixture of full-length FUS protein and RNA, including the dilute- and condensed-binodal branches as well as tie-lines (**Fig 5f**). We note that three pieces of information, tie-lines along with both binodals, are strictly required to physically relate the compositions of coexisting phases to the average composition specified in experiments, and that no other methods suitable for low-yield proteins provide all three. Further, we proved mathematically that the proposed methodology can, in principle, resolve the composition of condensates containing an arbitrary number of distinct molecular components (**Supplementary Note 5**). Whether partitioning of a particular species can be resolved is determined primarily by the sensitivity and precision of the dilute-phase detection method employed (**Extended Data Fig. 7**). We emphasize that our methodology is agnostic to the choice of dilute-phase detection method. This flexibility allows experimenters to select or combine established analytic approaches best-suited for the molecules used. In particular, mass-spectrometry can resolve complex mixtures with 100s or 1000s of components and distinguish between protein some PTM-states ^48^.

In future applications of this method, there are three classes of practical considerations that must be kept in mind. The first concerns condensate size. Though QPI measures refractive index robustly for droplets over the size-range most commonly encountered in reconstitution experiments, from a few to a few 10s of microns, systematic errors are incurred due to scattering and gravitational settling for sufficiently small and large droplets, respectively. We anticipate that additional development of fitting routines to account for these physical effects can further extend the size range for reliable measurements ^49,50^. The second consideration involves treating the refractive index of a mixture as a linear sum of contributions from its components (see **Eq. 3, Supplementary Note 2**). This is a very good approximation for protein solutions over a wide range ^30^, though higher-order terms could potentially contribute for some molecules ^51^. In the cases we checked, we found the linear sum to describe the refractive index of multi-component mixtures accurately to within a few % (**Extended Data Fig. 5**). We also note that while contributions from PTMs are currently neglected when estimating protein refractive index increments ^32^, which are the coefficients in the linear sum, this could be readily addressed in the future with molar refractivity measurements of modified amino acids in solution ^52^. This could improve accuracy for condensates containing highly-modified proteins. The third consideration is in regard to measurements in cells. Though our QPI-based method is best-suited for reconstituted condensates, QPI and related techniques like ODT can provide some information on cellular condensates whose refractive index differs sufficiently from the surrounding material ^53,31,35^. For condensates with sufficient contrast, reconstruction of binodals and tie-lines would require knowledge of molecular abundances in individual cells as well as the coexisting phase. Absent this, phase-based imaging data can still provide a quantitative measure of the average macromolecular mass difference between phases *in vivo*, which may be particularly informative in the context of perturbations.

As composition ultimately influences all other condensate properties and associated cellular functions, we anticipate a central role for this method in addressing many pressing biological questions. Measuring condensate composition as the abundance of individual components are systematically varied will reveal the thermodynamic contributions of these molecules to the phase, potentially clarifying the biological function of individual components. We anticipate that correlating composition with other physical properties like viscoelasticity, interfacial tension, and dielectric constant will likewise provide insight into condensate function. By providing a ground-truth with which to calibrate fluorophore behavior, we expect that this method will enable the use of dyes to both follow reactions localized to condensates as well as quantitatively probe the chemical environment within condensates. The latter will likely be essential to understand and potentially tune the partitioning of therapeutic drugs into condensates in treatment of diseases like cancer ^54^.

## MATERIALS AND METHODS

### Sample preparation

Recombinant protein constructs used in this work were purified and stored as described previously ^6,7^. To induce phase separation, we mixed protein in high-salt storage buffer (300 mM KCl for PGL3 constructs, 500 mM KCl for TAF15 and FUS constructs) with storage buffer lacking monovalent salt, “Dilution Buffer”, to reach the desired final salt concentration. Generally, an aliquot of Dilution Buffer was supplemented to 1 mM with fresh DTT prior to each day’s experiments. After induction of phase separation, dilute phase was obtained by centrifugation at RCF = 20,800 x g for 30 min in a tabletop centrifuge (Centrifuge 5417 R, Eppendorf) pre-equilibrated at the desired temperature. For control measurements, 10-µm silica microspheres were purchased from Whitehouse Scientific (Waverton, UK), and glycerol-water-mixtures were prepared by weight to the desired refractive index. Bead-containing dispersions were prepared by gently dipping a 10-µL pipette tip into a stock of dry beads, transferring the pipette tip to a 40 µL volume of glycerol-water mixture, and pipette mixing to disperse. Aqueous two-phase systems with PEG-35k (Sigma) and Dextran T500 (Pharmocosmos) were prepared as described previously ^34^. BSA was purchased from Sigma and used without further purification.

### Quantitative phase imaging and analysis

QPI measurements were performed using a coherence-controlled digital holographic microscope (Q-Phase, Telight (formerly TESCAN), Brno, CZ) based on the set-up in ^28^. Most data were acquired on a Generation-1 instrument with a tungsten-halogen bulb as lightsource, though some data were acquired on a Generation-2 instrument with a 660-nm LED as lightsource. In each case, the holography lightsource was filtered by a 10-nm bandwidth notch filter centered at 650 nm. All measurements were performed with 40x dry objectives (0.9 NA, Nikon) except those for SNAP-TAF15(RBD) reported in **Fig. 3f**, for which 20x dry objectives were used. In all cases, the condenser aperture was set to an NA of 0.30. Immediately following phase separation, ∼ 5 µL of sample was loaded into a temperature-controlled flowcell, sealed with two-component silicone glue Twinsil (Picodent, Wipperfürth, DE), and allowed to settle under gravity for ∼ 10 minutes prior to data collection. Flowcells were constructed with a 30×24×0.17 mm^3^ PEGylated coverslip and a 75×25×1 mm^3^ sapphire slide as bottom and top surfaces, respectively, using parafilm strips as spacers. Proportional-integral-derivative (PID)-controlled Peltier elements affixed to the sapphire slide enabled regulation of flowcell temperature, as previously described ^39^. The sapphire, coverslip, and spacers were adhered by heating the assembled flowcell to 50 °C for 2-5 min, then returning to the desired temperature for the first measurement, typically to 20 °C. For each sample, hologram *z*-stacks (*dz* = 0.2 µm, first plane typically near the coverglass surface) were acquired for several fields of view. SophiQ software (Telight, Brno, CZ) was used to construct amplitude and compensated phase images from the raw holograms. Pixels in the phase images are 0.157 µm per side for the 40x, and pixel intensities are in units of radians. To aid interpretation by persons with red/green color perception deficiencies, phase images are displayed using the Ametrine colormap ^55^.

All phase images were analyzed in MATLAB using custom code. For each *z*-plane, compensated phase images were first segmented to identify individual droplets. To determine the background phase value, *φ*_0_, the image’s pixel intensity histogram was fit to a Gaussian, and the Gaussian center taken as *φ*_0_. Pixel intensities *φ* ≥ *n*_*sig*_σ_*φ*_ are considered above threshold, where σ_*φ*_ is the standard deviation extracted from the Gaussian fit. Typically, *n*_*sig*_ = 5. A binary mask was generated with this threshold and individual objects were identified using the MATLAB function bwconncomp.m. For each object, a region of interest slightly larger than the object’s bounding box was fit twice to phase functions of the form given by Eq. (1) in the main text. First, we fit using the projected height of a sphere,

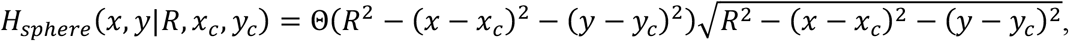

in order to obtain estimates for the parameters ∆*n, R, x*_*c*_, *y*_*c*_, where Θ(*x*) is the Heaviside function. These estimates were then used to initialize a fit to a regularized version of Eq. (1),

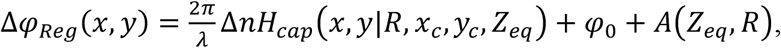

using the projected height of a spherical cap,

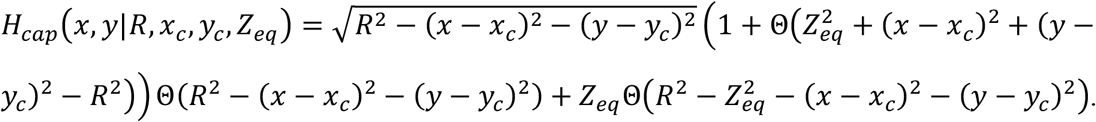

The regularization is given by *A*(*Z*_*eq*_, *R*) = *A*_0_ (*Z*_*eq*_ − *R*)_2_ Θ (*Z*_*eq*_ − *R*) with *A*_0_ = 10^5^ and *φ*_0_ fixed at the value obtained from the pixel intensity histogram fit. After all z-planes were processed, the objects were tracked through *z* using track.m (https://site.physics.georgetown.edu/matlab/index.html). For each tracked object, the representative fit parameters are taken as those from the fit with the highest Adj. R^2^ value, which are typically in the plane acquired nearest to the equatorial plane of a given droplet. The particle list was then automatically filtered for fit quality (typically retaining only Adj. R^2^-values > 0.95) and overlap with dead-pixels on the detector. Droplets with irregular wetting or that are not isolated in *z* (i.e. are situated beneath other droplets in solution) are removed manually following inspection of the raw data.

### Optical Diffraction Tomography

ODT measurements were performed using a custom-built microscope employing a 532-nm laser, as described previously ^56^. Tomogram reconstruction and image analysis was performed as described previously ^57,10^.

### Confocal Fluorescence Microscopy and Analysis

Confocal imaging was performed on an inverted Zeiss LSM880 point-scanning confocal microscope with a 40x water-immersion objective (1.2 NA, C-Apochromat, Zeiss) at room temperature. mEGFP was excited with a 488-nm argon laser and emission detected with a 32-channel GaAsP photomultiplier tube (PMT) array set to accept photon wavelengths between 499 and 569 nm. AlexaFluor546 was excited with a 561-nm diode-pumped solid-state laser and emission detected between 570 and 624 nm with the same spectral PMT array. For both fluorophores, the confocal pinhole diameter was set to 39.4 µm, corresponding to 0.87 and 0.96 Airy Units for mEGFP and AlexaFluor546, respectively. For each field of view, scanning was performed with a lateral pixel size of 0.415 µm and *z*-stacks acquired with a spacing of 0.482 µm.

All confocal fluorescence images were analyzed in MATLAB using custom code. Partition coefficients of fluorescently-labeled species into condensates are estimated on the basis of the fluorescence intensity along a line-scan through the droplet center. The analysis pipeline begins with determining the location of each condensate and an appropriate line-scan orientation angle. To determine lateral positions of condensates in each field of view, a *z*-plane was selected slightly above the coverglass such that even small droplets appeared bright. Following convolution with a 2D Gaussian (σ_*x*_ = σ_*y*_ = 0.5 pixels) to suppress shot noise, a threshold of *I*_*thresh*_ = max(*I*(*x, y*)) /2 was applied to obtain a binary mask. The lateral positions and approximate sizes of objects were determined from the mask with bwconncomp.m. Only the largest ∼120 objects for each condition were analyzed further. For each object, partition coefficients were calculated using the z-plane for which the mean intensity in a 5-pixel-radius disk concentric with the object was greatest. Line-scans were 51 pixels long, concentric with the object, and averaged over a width of 3 pixels. Suitable line-scan orientations were determined in a semi-automated manner by superimposing reference lines rotated through 15° increments on each object and manually selecting an orientation that best avoided neighboring objects. Objects for which no suitable line-scan orientation could be found were discarded.

Line-scans for each droplet were automatically subdivided into three domains, corresponding to pixels in the dilute phase, the condensed phase, or the exclusion zone. The positions of the left and right dilute/condensed interfaces are estimated as those at which the intensity profile reaches its half-maximal value above detector background *I*_*BKgd*_ (see below). To reduce artefacts stemming from the finite point-spread function of the microscope, pixels within the greater of 1 pixel or *l*_*EZ*_ = 1.22λ/(2*NA*) on either size of the half-maximum were excluded from the analysis. Remaining profile pixels outside the droplet were averaged to give *I*_*Dil*_, while profile pixels inside are averaged to give *I*_*Cond*_. The partition coefficient for each object was calculated according to

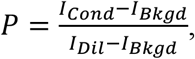

where *I*_*BKgd*_ is the average of all pixels in a background image acquired immediately following the fluorescence *z*-stack. Background images were acquired with the light source blocked to measure the contribution of detector noise to the signal.

### Bulk Refractometry

Data in Fig. 2i were acquired at λ = 653.3 nm and 21 °C with a DSR-L multi-wavelength refractometer (Schmidt + Haensch, Berlin) using a 200-µL sample volume. All other bulk refractive index measurements were acquired at λ = 589.3 nm with a J457 refractometer (Rudolph Research Analytical, Hackettstown, NJ). The refractive index of glycerol/water mixtures was adjusted from λ = 589.3 to 650 nm using empirical dispersion relations for distilled water, *n*_*Water*_(λ) ^58^, and glycerol, *n*_*Glycerol*_(λ) ^59^ according to

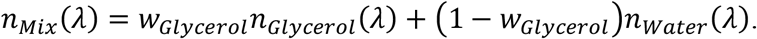

The glycerol weight fraction in the mixture was calculated from the refractive index measurement of the mixture at 589.3 nm as

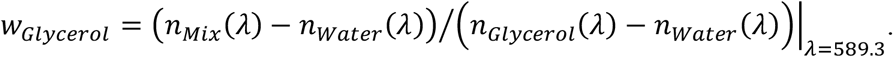

### Bead porosity models

The two models used to account for the porosity *p* of the silica microspheres (Fig. 2h, Extended Data Fig. 2) are a weighted linear sum, Δ*n* = *pn*_*Silica*_ + (1 − *p*)*n*_*Water*_ (simple model), and a more detailed model,

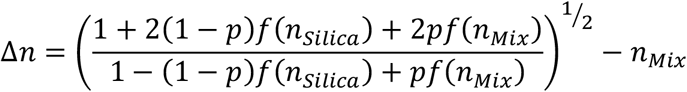

based on the Lorentz-Lorenz relation ^60^, wherein

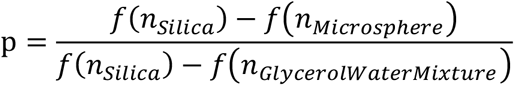

with *f*(*n*) ≡ (*n*^2^ − 1)/(*n*^2^ + 2).

### Calculation of *dn*/d*c*, 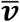 and polymer volume fraction

The refractive index increment and partial specific volume were estimated for each protein construct using the calculator tool within SEDFIT ^32^ and the protein sequences listed in the SI. The partial specific volume of 0.5773 mL/g for poly(A) RNA was estimated using consensus volumes per base from ^61^ and assuming a typical chain length of 500 bases. The partial specific volumes of 0.8321 and 0.6374 mL/g for PEG-35k and Dextran-500k, respectively, were taken from ^34^. Polymer volume fraction (Fig. 5f) for polymer *i* is given by 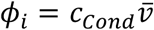

### Calculations for aging systems

Volumes of individual aging condensates (Fig. 4c,f) were calculated as 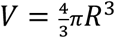 The number of molecules in each condensate (Fig. 4d,g) were calculated as *N*(*t*) = *c*(*t*)*V*(*t*). Given *c*(*t*), the relative volume change expected if *N*(*t*) = *N*(0) is given by *V*(*t*)/*c*(0) = *c*(0)/*c*(*t*) (Fig. 4f, black line).

### UV-Vis spectroscopy

Absorption spectra of dilute-phase and reference samples for the FUS/RNA system (Fig. 5) were collected on an NP-80 spectrophotometer (IMPLEN, München). All spectra were acquired at room temperature over λ ∈ [200 nm, 900 nm]. For each raw spectra 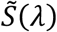 a linear fit on λ ∈ [550 nm, 750 nm] was used to determine a baseline correction, *S*_*BL*_ (λ). Corrected spectra are given by 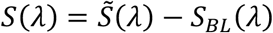 At least 3 replicate spectra were acquired for each condition and averaged following baseline correction to give the final representative spectra. The uncertainty in the spectra at each wavelength was estimated as the standard deviation of the corrected replicates. Dilute-phase spectra were demixed (Fig. 5b) into a weighted sum of three contributions

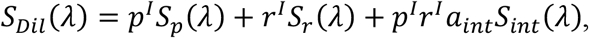

where (*p*^*I*^, *r*^*I*^) are the protein and RNA concentrations in the dilute phase. *S*_*p*_ and *S*_*r*_ are reference spectra for protein and RNA, respectively. The final term captures the effect of protein-RNA interactions on the absorbance of a mixture, which could physically stem from binding-induced changes in extinction coefficients. Reference spectra for the interaction was calculated from spectra of a protein/RNA mixture of known composition (*p, r*) in the 1-phase regime according to *S*_*int*_ = *S*(*p, r*) − *pS*_*p*_ − *rS*_*r*_. The parameter α^*int*^ captures the approximately linear increase of *Si*^*nt*^ with p. The same value of α_*int*_ was used to demix all dilute-phase spectra. The dilute-phase concentrations *p*^*I*^, *r*^*I*^ were therefore the only free parameters for the demixing.

## ACKNOWLEDGEMENTS

We thank E. Filippidi, T. Franzmann, B. Diederich, T. Harmon, L. Jawerth, A.W. Fritsche, and J.M. Iglesias-Artola, as well as members of the Brugués, Hyman, Alberti, Jülicher, and Weber groups for helpful discussions. We thank T. Slabý, M. Šicner, and V. Procházka of Telight for support with the Q-Phase microscope and E. Bittrich and R. Schulze of the Leibniz Institute for Polymer Research for assistance with multi-wavelength refractometry measurements. We thank A. Poznyakovskiy for creating several constructs for this project, J. Wang for purification of TAF15 protein constructs, R. Lemaitre for purification of full-length FUS protein, D. Kuster for the gift of poly(A) RNA, and A.W. Fritsche for the gift of silica beads. We thank A. Schwager and S. Ernst for preparation of passivated coverglass, L. Jawerth for assistance with aging PGL3 samples, and A.W. Fritsche for assistance with the temperature stage. We also thank the MPI-CBG Core Facilities, particularly the Light Microscopy Facility and the Protein Expression and Purification Core, for their support. PMM was supported by an ELBE Postdoctoral Fellowship from the CSBD. PMM and JB were also supported by Volkswagen ‘Life’ grant number 96827.

## AUTHOR CONTRIBUTIONS

PMM, JP, AAH, and JB conceived the project. PMM conceived and developed the multi-component measurement approach and derived the analytical solution. PMM designed and performed QPI, refractometry, spectroscopy, and fluorescence microscopy measurements, developed the analysis pipelines for these data types, and performed the formal analysis. KK and PMM designed and performed ODT measurements. KK analyzed ODT data. MR-G purified multiple constructs and articulated essential distinctions related to the handling of IDRs and multi-domain proteins. JB, AAH, and JG supervised the work. PMM and JB wrote the paper with input from all authors.

## DATA AVAILABILITY

Data generated and analyzed supporting the findings of this Article will be made available by the Authors upon reasonable request.

## CODE AVAILABILITY

Custom code generated supporting the findings of this manuscript will be made available by the Authors upon reasonable request.

## SUPPLEMENTARY INFORMATION

- Note 1: Requirements of bulk measurements
- Note 2: Linear approximation to refractive index of mixtures
- Note 3: Measurement of c_Dil_
- Note 4: Requirement that tie-lines are straight
- Note 5: Derivation of condensate composition for multi-component systems
- Supplementary
- References
- Table S1: Protein constructs used
- List S1: List of protein sequences used

## EXTENDED DATA FIGURES AND CAPTIONS

**Extended Data Fig. 1:**
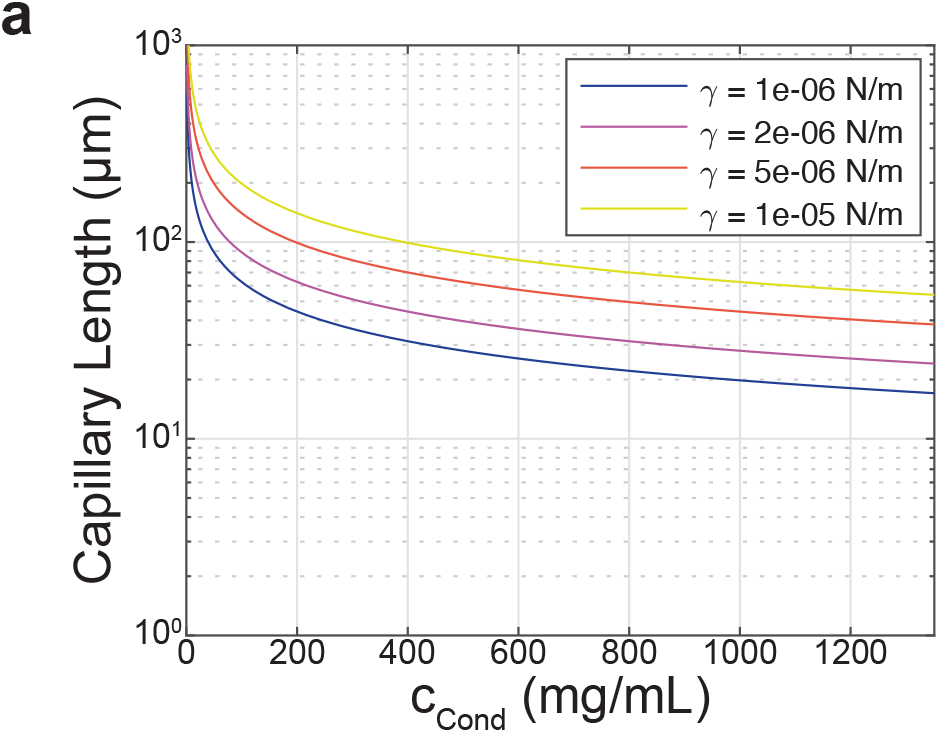
Estimate of capillary length for biomolecular condensates. **a**, Capillary length as a function of the condensed-phase polymer mass concentration. The capillary length increases with increasing interfacial tension, so we plot traces for different values of interfacial tension. From bottom to top, interfacial tensions are 1, 2, 5, and 10 µN/m. This spans the range of tensions reported for PGL-3 condensates by ^62^. For a given droplet density and interfacial tension, the spherical cap approximation for the droplet shape is valid so long as the droplet size is less than the capillary length. In the case of PGL-3, our density measurements of ∼ 90 mg/ml indicate a capillary length of 67 µm at the lowest interfacial tension, which is much larger than the 1-8 µm radii of the droplets. Although the capillary length is reduced to ∼ 30 µm for the higher ∼ 400-500 mg/ml densities we measure for TAF15(RBD) condensates, this length is still much larger than the 0.67-3.2 µm radii of those condensates analyzed in this work.

**Extended Data Fig. 2:**
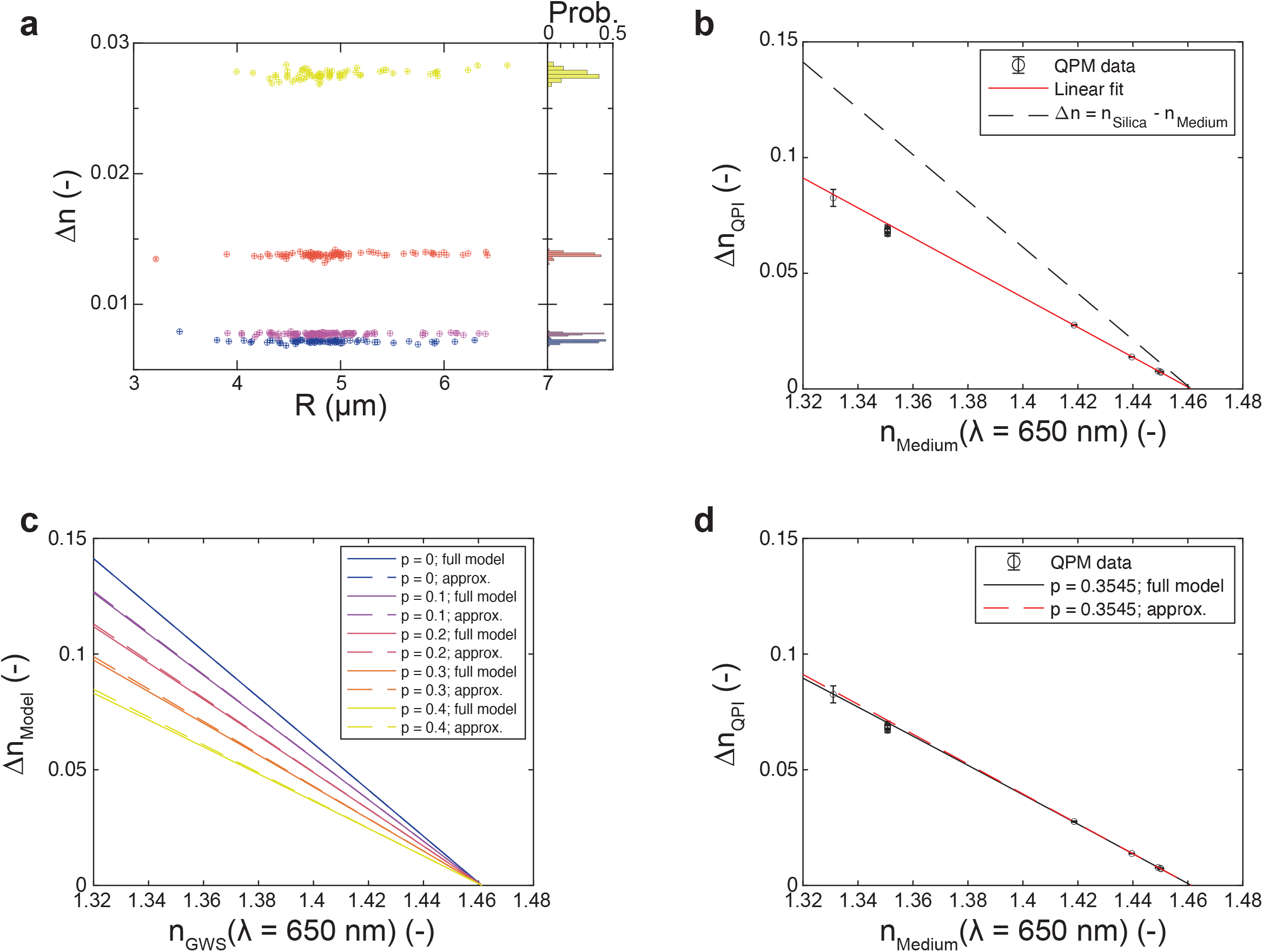
Silica bead measurements and porosity. **a**, (left) Refractive index difference between silica beads and the surrounding glycerol/water mixture as a function of bead size as measured by QPI for a range of glycerol/water ratios. Glycerol weight fraction of the medium is 0.6641 (yellow, *N* = 66), 0.808 (orange, *N* = 83), 0.8688 (purple, *N* = 102), and 0.8751 (blue, *N* = 74). Datapoints represent individual beads, error bars are 95% confidence intervals returned from fits, and are typically smaller than the datapoint. Though the nominal bead diameter is 10 µm, the samples show significant polydispersity. Importantly, there is no strong systematic variation of the extracted ∆*n* with bead size. (right) Measured refractive index distribution is approximately symmetric for each condition. **b**, Population mean (± standard deviation) of ∆*n* measured by QPI at λ = 650 nm for silica beads as a function of the refractive index of the glycerol/water mixture, *n*_*medium*_. Note that *n*_*medium*_ was measured at λ = 589 nm on a digital refractometer and is plotted following adjustment to λ = 650 nm using published dispersion relationships for glycerol and distilled water and an approximate mixing rule, as described in Methods. Solid red line is a linear fit to the data with the 4 largest *x*-values, with slope -0.6455 ± 0.01806. The *x*-intercept of the fit line, at which the refractive index of the silica bead is indistinguishable from that of the medium, is 1.461 ± 0.057. Importantly, this is comparable to published values of fused silica under similar conditions (e.g. n = 1.4565 at 650 nm, 20 °C, ^63^), independently of whether we adjust for dispersion. However, the slope of the fit line (−0.6455 ± 0.0181) differs significantly from the value of -1 expected for beads of pure fused silica (dashed black line). This suggests that the beads may be porous, as has been reported previously ^60^. **c**, The refractive index difference expected for silica beads with porosity *p* as a function of the refractive index of the surrounding glycerol/water mixture, assuming that the bead pores are filled with the glycerol/water mixture. Here *p* is the fraction of the bead volume occupied by pores rather than silica. Different colors represent different values of porosity from 0 to 0.4. For each *p*, ∆*n* is calculated using two different refractive index mixing rules, either a weighted linear combination (approx., dashed line) or the Lorentz-Lorenz relation (full model, solid line, Methods). Both models predict that fluid-filled pores in the silica beads would give a ∆*n* that decreases (nearly) linearly with slope (1-*p*) as n_medium_. increases. The two models are almost indistinguishable except at low *n*_*medium*_ and high *p*, where the simple approximation noticeably overestimates ∆*n*. **d**, Same data as in **b** above, but now overlaid with model predictions for silica beads with *p* = 0.3545. All measurements at 25 °C.

**Extended Data Fig. 3:**
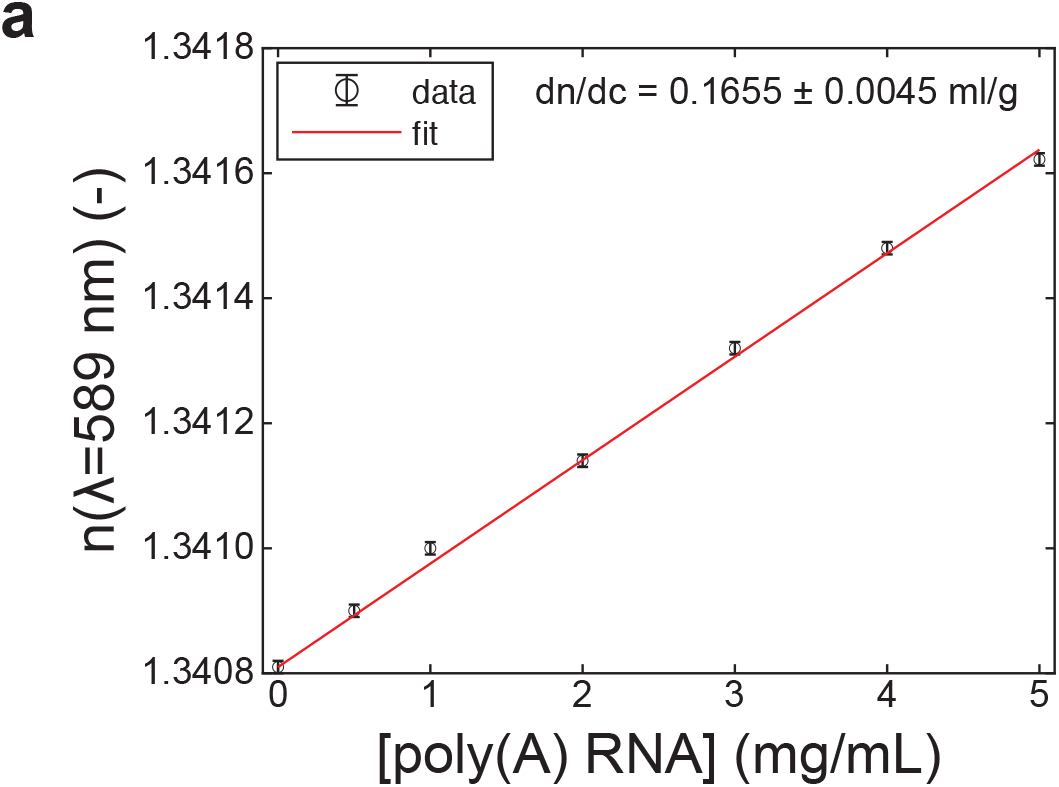
Measurement of dn/dc for poly(A) RNA at 589 nm. **a**, Refractive index of poly(A) RNA in water as a function of RNA concentration. Measurements were performed with a digital refractometer at 589 nm at 20 °C. Error bars are the larger of the standard deviation of *N* = 5 measurements or the instrument resolution 0.00001 (if all repeat measurements were identical). The refractive index increment is given by a linear fit as 0.1655 ± 0.0045 ml/g.

**Extended Data Fig. 4:**
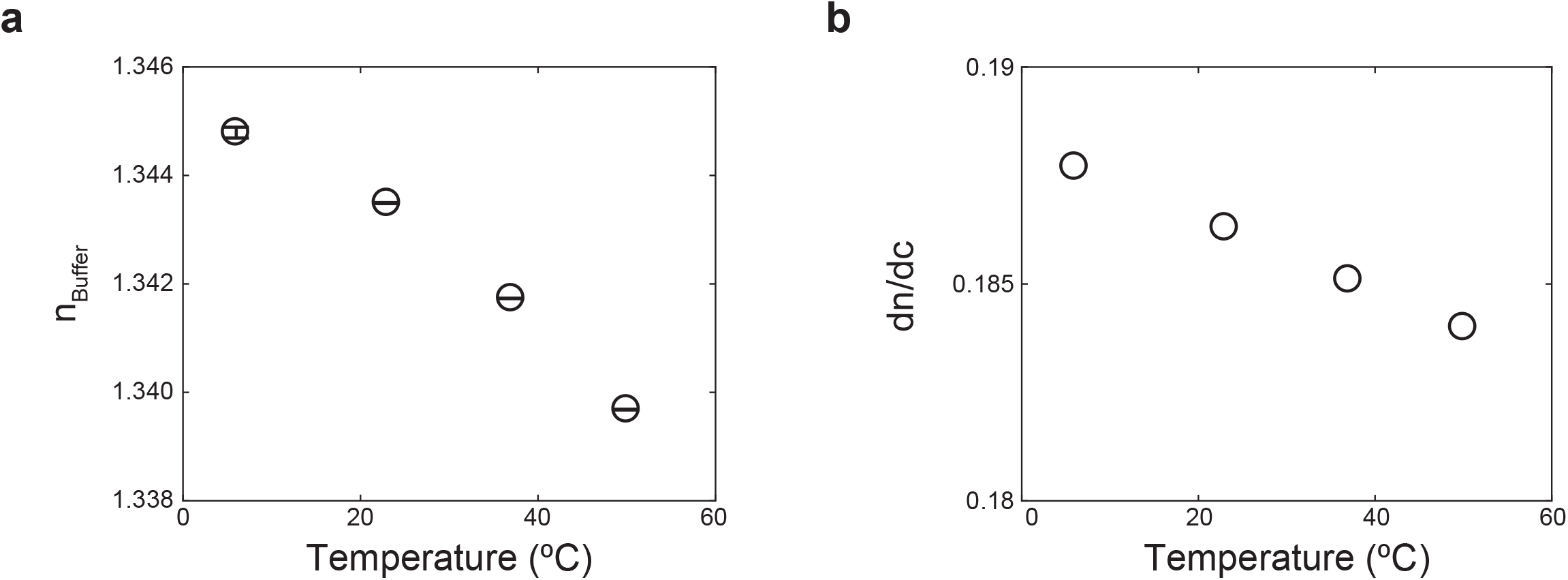
Temperature-dependent optical constants. **a**, Refractive index of buffer as measured with a digital refractometer at 589 nm at the temperatures used in Fig. 3f. Error bars are the larger of the standard deviation of *N* = 5 measurements or the instrument resolution 0.00001 (if all repeat measurements were identical). The refractive index of the solution decreases with increasing temperature primarily due to the reduced electron density accompanying thermal expansion. Note that the reduction in refractive index due to this effect, *n*_*buffer*_(6 °C) - *n*_*buffer*_(50 °C) = 0.0051, is much smaller than the variation of ∆*n* we measure for TAF15(RBD) condensates over the same range, ∆*n*(6 °C) - ∆*n*(50 °C) = 0.0144. **b**, Refractive index increment for SNAP-TAF15(RBD) estimated using SEDFIT software ^32^ as a function of temperature. These values were used in Fig. 3f to convert the ∆*n* measured by QPI to the *c*_*cond*_ values reported at each temperature.

**Extended Data Fig. 5:**
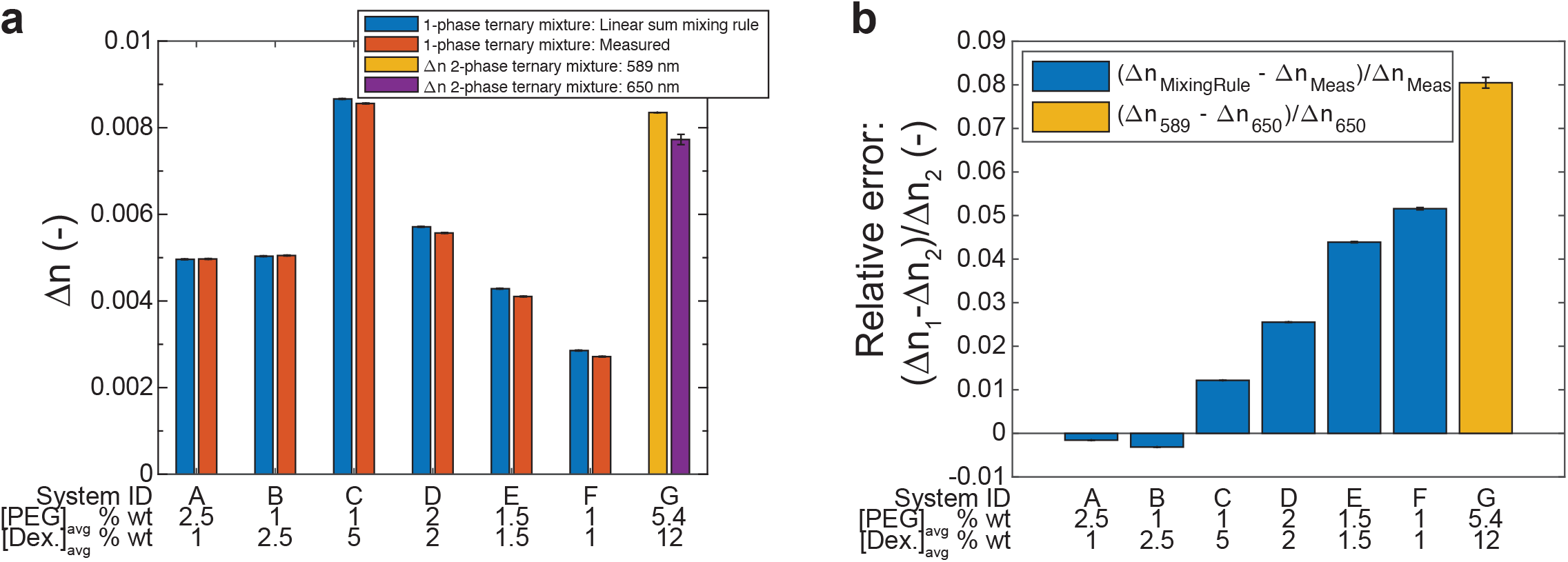
Validation of linear sum approximation for refractive index of mixtures. **a**, Refractive index difference between either homogeneous PEG/Dextran mixtures prepared in the one-phase regime and water (blue and red) or between coexisting phases in the two-phase regime (yellow and purple). Average system composition is specified below each bar as weight fractions of PEG-35k and Dextran-500k. Blue bars show the refractive index measured experimentally on a digital refractometer at 589 nm, while red bars show the refractive index predicted for the same compositions using a linear sum approximation for the mixing rule (see Methods). Yellow bar shows the difference between the refractive indices of the coexisting phases, each measured individually at 589 nm on a digital refractometer. Purple bar shows the population mean refractive index difference between the same phases measured via QPI for *N* = 205 droplets. Error bars for digital refractometry measurements are the larger of the standard deviation of *N* = 5 measurements or the instrument resolution 0.00001 (if all repeat measurements were identical). Error bars for the predictions are estimated at 0.00001. Error bar for the QPI measurement is the standard deviation of the measured ∆*n* distribution. All measurements were performed at 24 °C. **b**, Relative error of the paired measurements in **a**. The relative error incurred by application of the linear sum mixing rule is typically much less than 5 %. For comparison, natural variation in optical properties with wavelength (dispersion) results in an 8% change in ∆*n* between 589 nm and 650 nm.

**Extended Data Fig. 6:**
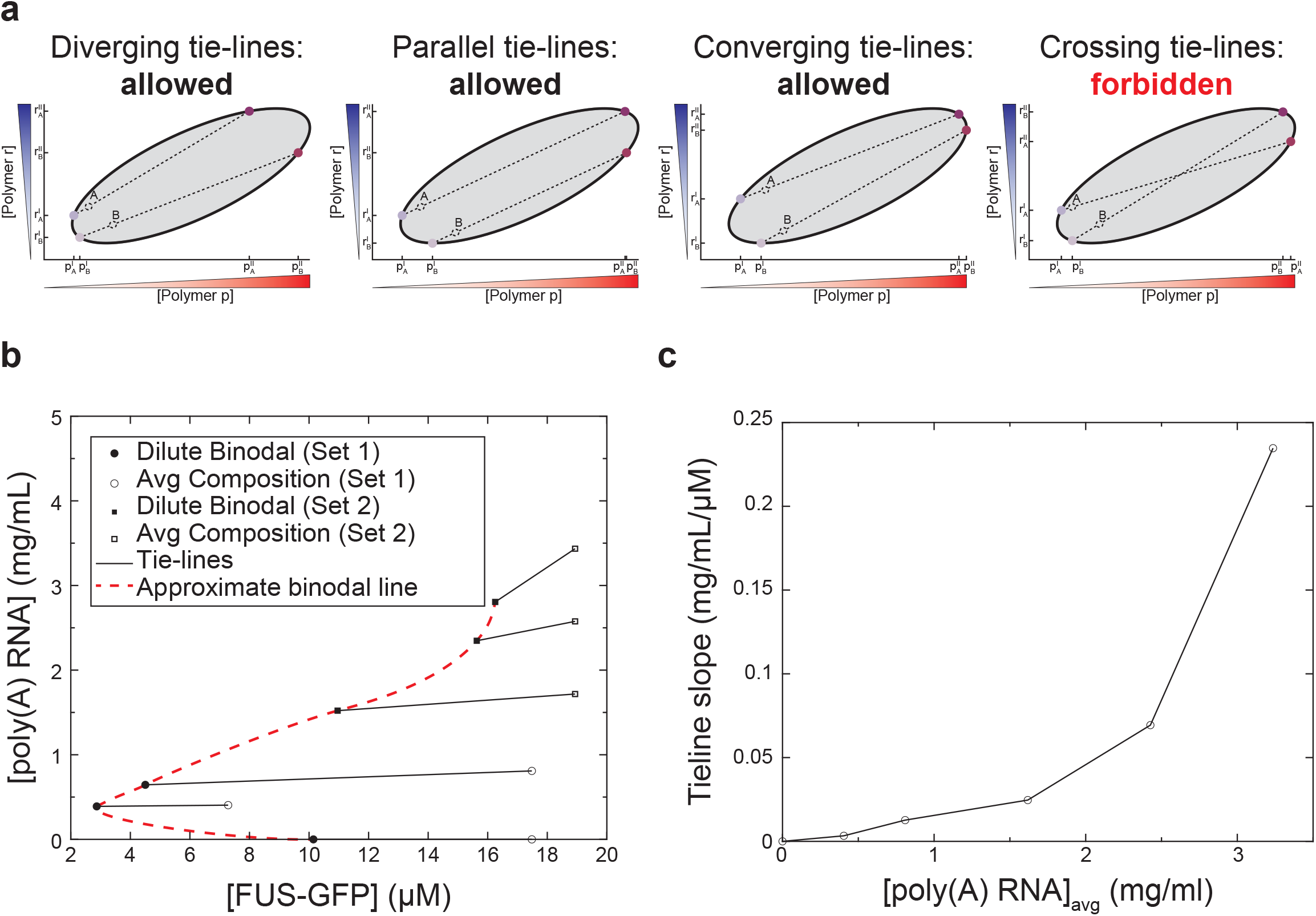
Examples of physically permissible tie-lines. **a**, Examples of ternary phase diagrams with varying tie-line orientations. While tie-lines can in principle diverge (left), be parallel (center-left), or converge (center-right), it is unphysical and thus forbidden for them to cross within the multi-phase coexistence region (right). **b**, Ternary phase diagram for FUS/poly(A) RNA on linear scales zoomed in on the dilute binodal. **c**, Slopes of the tie-lines shown in **b**. Relative to the classification scheme introduced in **a**, this system displays diverging tie-lines.

**Extended Data Fig. 7:**
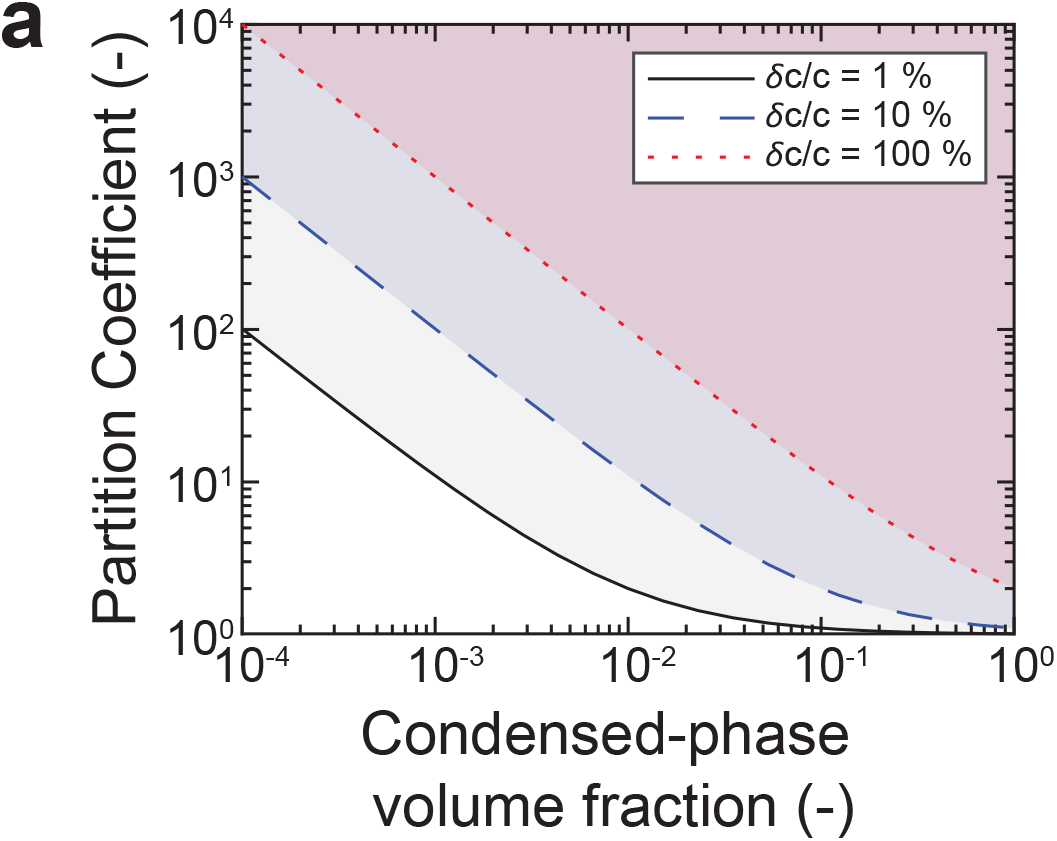
Multi-component detection limits. **a**, The minimum partition coefficient P_*min*_ for which the concentration of a species in the dilute phase *c*^*I*^ is measurably different from its total average concentration in the system 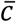 depends on the relative precision *δc*/*c* of the dilute-phase detection strategy. For a given *δc*/*c, P*_*min*_ decreases as the volume fraction of the condensed phase increases. In this diagram, curves show the minimum partition coefficient for which partitioning of an arbitrary species would be detectable in a system as a function of condensed-phase volume fraction for *δc*/*c* = 1% (solid black), 10% (blue dashed), and 100 % (red dotted). The shading denotes that all partition coefficients above the corresponding lower-limit are detectable.

### Supplementary Video 1

Quantitative phase imaging of aging timecourse for reconstituted PGL3 condensates.

## REFERENCES

1. Hyman, A. A., Weber, C. A. & Julicher, F. Liquid-Liquid Phase Separation in Biology. Annu. Rev. Cell Dev. Biol. 30, 39–58 (2014).

2. Banani, S. F., Lee, H. O., Hyman, A. A. & Rosen, M. K. Biomolecular condensates: organizers of cellular biochemistry. Nat. Rev. Mol. Cell Biol. 18, 285–298 (2017).

3. Alberti, S., Gladfelter, A. & Mittag, T. Considerations and Challenges in Studying Liquid-Liquid Phase Separation and Biomolecular Condensates. Cell 176, 419–434 (2019).

4. Elbaum-Garfinkle, S. et al. The disordered P granule protein LAF-1 drives phase separation into droplets with tunable viscosity and dynamics. Proc. Natl. Acad. Sci. 112, 7189–7194 (2015).

5. Nott, T. J. et al. Phase Transition of a Disordered Nuage Protein Generates Environmentally Responsive Membraneless Organelles. Mol. Cell 57, 936–947 (2015).

6. Saha, S. et al. Polar Positioning of Phase-Separated Liquid Compartments in Cells Regulated by an mRNA Competition Mechanism. Cell 166, 1572-1584.e16 (2016).

7. Wang, J. et al. A Molecular Grammar Governing the Driving Forces for Phase Separation of Prion-like RNA Binding Proteins. Cell 174, 688-699.e16 (2018).

8. Updike, D. & Strome, S. P Granule Assembly and Function in Caenorhabditis elegans Germ Cells. J. Androl. 31, 53–60 (2010).

9. Xing, W., Muhlrad, D., Parker, R. & Rosen, M. K. A quantitative inventory of yeast P body proteins reveals principles of composition and specificity. eLife 9, e56525 (2020).

10. Guillén-Boixet, J. et al. RNA-Induced Conformational Switching and Clustering of G3BP Drive Stress Granule Assembly by Condensation. Cell 181, 346-361.e17 (2020).

11. Riback, J. A. et al. Composition-dependent thermodynamics of intracellular phase separation. Nature 581, 209–214 (2020).

12. Kaur, T. et al. Sequence-encoded and composition-dependent protein-RNA interactions control multiphasic condensate morphologies. Nat. Commun. 12, 872 (2021).

13. Alshareedah, I., Thurston, G. M. & Banerjee, P. R. Quantifying viscosity and surface tension of multicomponent protein-nucleic acid condensates. Biophys. J. 120, 1161– 1169 (2021).

14. Banani, S. F. et al. Compositional Control of Phase-Separated Cellular Bodies. Cell 166, 651–663 (2016).

15. Choi, S., Meyer, M. O., Bevilacqua, P. C. & Keating, C. D. Phase-specific RNA accumulation and duplex thermodynamics in multiphase coacervate models for membraneless organelles. Nat. Chem. 1–8 (2022) doi:10.1038/s41557-022-00980-7.

16. Morin, J. A. et al. Sequence-dependent surface condensation of a pioneer transcription factor on DNA. Nat. Phys. 1–6 (2022) doi:10.1038/s41567-021-01462-2.

17. Lakowicz, J. R. Solvent and Environmental Effects. in Principles of Fluorescence Spectroscopy 205–235 (Springer, Boston, MA, 2006). doi:10.1007/978-0-387-46312-4_6.

18. Küffner, A. M. et al. Acceleration of an Enzymatic Reaction in Liquid Phase Separated Compartments Based on Intrinsically Disordered Protein Domains. ChemSystemsChem 2, e2000001 (2020).

19. Avni, A., Joshi, A., Walimbe, A., Pattanashetty, S. G. & Mukhopadhyay, S. Single-droplet surface-enhanced Raman scattering decodes the molecular determinants of liquid-liquid phase separation. Nat. Commun. 13, 4378 (2022).

20. Broide, M. L., Berland, C. R., Pande, J., Ogun, O. O. & Benedek, G. B. Binary-liquid phase separation of lens protein solutions. Proc. Natl. Acad. Sci. 88, 5660–5664 (1991).

21. Zhang, F. et al. Charge-controlled metastable liquid–liquid phase separation in protein solutions as a universal pathway towards crystallization. Soft Matter 8, 1313–1316 (2012).

22. Li, L. et al. Phase Behavior and Salt Partitioning in Polyelectrolyte Complex Coacervates. Macromolecules (2018) doi:10.1021/acs.macromol.8b00238.

23. Brady, J. P. et al. Structural and hydrodynamic properties of an intrinsically disordered region of a germ cell-specific protein on phase separation. Proc. Natl. Acad. Sci. 114, E8194–E8203 (2017).

24. Martin, E. W. et al. Valence and patterning of aromatic residues determine the phase behavior of prion-like domains. Science 367, 694–699 (2020).

25. Bremer, A. et al. Deciphering how naturally occurring sequence features impact the phase behaviors of disordered prion-like domains. 2021.01.01.425046 https://www.biorxiv.org/content/10.1101/2021.01.01.425046v1 (2021) doi:10.1101/2021.01.01.425046.

26. Jawerth, L. et al. Protein condensates as aging Maxwell fluids. Science 370, 1317–1323 (2020).

27. Park, Y., Depeursinge, C. & Popescu, G. Quantitative phase imaging in biomedicine. Nat. Photonics 12, 578–589 (2018).

28. Slabý, T. et al. Off-axis setup taking full advantage of incoherent illumination in coherence-controlled holographic microscope. Opt. Express 21, 14747–14762 (2013).

29. Born, M. & Wolf, E. Principles of Optics. (Cambridge University Press, 2019).

30. Barer, R. & Tkaczyk, S. Refractive Index of Concentrated Protein Solutions. Nature 173, 821–822 (1954).

31. Biswas, A., Kim, K., Cojoc, G., Guck, J. & Reber, S. The Xenopus spindle is as dense as the surrounding cytoplasm. Dev. Cell 56, 967-975.e5 (2021).

32. Zhao, H., Brown, P. H. & Schuck, P. On the Distribution of Protein Refractive Index Increments. Biophys. J. 100, 2309–2317 (2011).

33. Park, J., Park, J., Lim, H. & Kim, H.-Y. Shape of a large drop on a rough hydrophobic surface. Phys. Fluids 25, 022102 (2013).

34. Atefi, E., Fyffe, D., Kaylan, K. B. & Tavana, H. Characterization of Aqueous Two-Phase Systems from Volume and Density Measurements. J. Chem. Eng. Data 61, 1531–1539 (2016).

35. Schlüßler, R. et al. Correlative all-optical quantification of mass density and mechanics of subcellular compartments with fluorescence specificity. eLife 11, e68490 (2022).

36. Matthews, B. W. Solvent content of protein crystals. J. Mol. Biol. 33, 491–497 (1968).

37. Lakowicz, J. R. Quenching of Fluorescence. in Principles of Fluorescence Spectroscopy 277–330 (Springer, Boston, MA, 2006). doi:10.1007/978-0-387-46312-4_8.

38. Iserman, C. et al. Condensation of Ded1p Promotes a Translational Switch from Housekeeping to Stress Protein Production. Cell 181, 818-831.e19 (2020).

39. Mittasch, M. et al. Non-invasive perturbations of intracellular flow reveal physical principles of cell organization. Nat. Cell Biol. 20, 344–351 (2018).

40. Fritsch, A. W. et al. Local thermodynamics govern formation and dissolution of Caenorhabditis elegans P granule condensates. Proc. Natl. Acad. Sci. 118, e2102772118 (2021).

41. Lin, Y., Protter, D. S. W., Rosen, M. K. & Parker, R. Formation and Maturation of Phase-Separated Liquid Droplets by RNA-Binding Proteins. Mol. Cell 60, 208–219 (2015).

42. Strom, A. R. et al. Phase separation drives heterochromatin domain formation. Nature advance online publication, (2017).

43. Shin, Y. & Brangwynne, C. P. Liquid phase condensation in cell physiology and disease. Science 357, eaaf4382 (2017).

44. Banerjee, P. R., Milin, A. N., Moosa, M. M., Onuchic, P. L. & Deniz, A. A. Reentrant Phase Transition Drives Dynamic Substructure Formation in Ribonucleoprotein Droplets. Angew. Chem. Int. Ed. 56, 11354–11359 (2017).

45. Maharana, S. et al. RNA buffers the phase separation behavior of prion-like RNA binding proteins. Science eaar7366 (2018) doi:10.1126/science.aar7366.

46. Choi, J.-M., Holehouse, A. S. & Pappu, R. V. Physical Principles Underlying the Complex Biology of Intracellular Phase Transitions. Annu. Rev. Biophys. 49, 107–133 (2020).

47. Lin, Y.-H., Brady, J. P., Chan, H. S. & Ghosh, K. A unified analytical theory of heteropolymers for sequence-specific phase behaviors of polyelectrolytes and polyampholytes. J. Chem. Phys. 152, 045102 (2020).

48. Valverde, J. M. et al. Single-embryo phosphoproteomics reveals the importance of intrinsic disorder in cell cycle dynamics. 2021.08.29.458076 https://www.biorxiv.org/content/10.1101/2021.08.29.458076v1 (2021) doi:10.1101/2021.08.29.458076.

49. Müller, P., Schürmann, M., Girardo, S., Cojoc, G. & Guck, J. Accurate evaluation of size and refractive index for spherical objects in quantitative phase imaging. Opt. Express 26, 10729–10743 (2018).

50. Müller, P., Cojoc, G. & Guck, J. DryMass: handling and analyzing quantitative phase microscopy images of spherical, cell-sized objects. BMC Bioinformatics 21, 226 (2020).

51. Barer, R. & Joseph, S. Refractometry of Living Cells: Part I. Basic Principles. J. Cell Sci. s3-95, 399–423 (1954).

52. McMeekin, T. L., Groves, M. L. & Hipp, N. J. Refractive Indices of Amino Acids, Proteins, and Related Substances. in Amino Acids and Serum Proteins vol. 44 54–66 (American Chemical Society, 1964).

53. Handwerger, K. E., Cordero, J. A. & Gall, J. G. Cajal Bodies, Nucleoli, and Speckles in the Xenopus Oocyte Nucleus Have a Low-Density, Sponge-like Structure. Mol. Biol. Cell 16, 202–211 (2005).

54. Klein, I. A. et al. Partitioning of cancer therapeutics in nuclear condensates. Science 368, 1386–1392 (2020).

55. Geissbuehler, M. & Lasser, T. How to display data by color schemes compatible with red-green color perception deficiencies. Opt. Express 21, 9862–9874 (2013).

56. Abuhattum, S. et al. Intracellular Mass Density Increase Is Accompanying but Not Sufficient for Stiffening and Growth Arrest of Yeast Cells. Front. Phys. 6, (2018).

57. Kim, K. et al. High-resolution three-dimensional imaging of red blood cells parasitized by Plasmodium falciparum and in situ hemozoin crystals using optical diffraction tomography. J. Biomed. Opt. 19, 011005 (2013).

58. Daimon, M. & Masumura, A. Measurement of the refractive index of distilled water from the near-infrared region to the ultraviolet region. Appl. Opt. 46, 3811–3820 (2007).

59. Rheims, J., Köser, J. & Wriedt, T. Refractive-index measurements in the near-IR using an Abbe refractometer. Meas. Sci. Technol. 8, 601–605 (1997).

60. Cheong, F. C., Xiao, K., Pine, D. J. & Grier, D. G. Holographic characterization of individual colloidal spheres’ porosities. Soft Matter 7, 6816–6819 (2011).

61. Voss, N. R. & Gerstein, M. Calculation of standard atomic volumes for RNA and comparison with proteins: RNA is packed more tightly. J. Mol. Biol. 346, 477–492 (2005).

62. Jawerth, L. M. et al. Salt-Dependent Rheology and Surface Tension of Protein Condensates Using Optical Traps. Phys. Rev. Lett. 121, 258101 (2018).

63. Malitson, I. H. Interspecimen Comparison of the Refractive Index of Fused Silica*,†. JOSA 55, 1205–1209 (1965).

