## Supplemental Information for "Label-free composition determination for biomolecular condensates with an arbitrarily large number of components"

### Supplementary information related to "Label-free composition determination of biomolecular condensates with an arbitrary number of components"

P.M. McCall et al.

#### Contents

**1 Note 1: Requirements of bulk measurements** **1**
1.1 Measurements above $c_{Cond}$ . . . . . . 1
1.2 Measurements below $c_{Dil}$ . . . . . . 2

**2 Note 2: Linear approximation to refractive index of mixtures** **2**

**3 Note 3: Measurement of $c_{Dil}$** **3**

**4 Note 4: Requirement that tie-lines are straight** **3**

**5 Note 5: Derivation of condensate composition for multi-component systems** **4**

**6 Bibliography** **11**

**7 Protein sequences used** **11**

### 1 Note 1: Requirements of bulk measurements

Ideally, one would measure $\frac{dn}{dc}$ and validate linearity over the entire range of $c \in [0, c_{Cond}]$. However, for proteins that phase separate, a large subset of this range lies in the miscibility gap, $c \in (c_{Dil}, c_{Cond})$, and so solutions with these concentrations *cannot be prepared*, at least not at the desired salt, pH, temp, etc. This leaves the formal possibility of comparing $\frac{dn}{dc}$ on $c \in [0, c_{Dil}]$ or for $c > c_{Cond}$. We first discuss issues for measurements above $c_{Cond}$, and subsequently for below $c_{Dil}$.

#### 1.1 Measurements above $c_{Cond}$

Excessive material requirements make the high concentration range impractical for many purified full-length proteins. To see this, we estimate the mass of purified protein required for a potential measurement.

To prepare samples of volume $V_{sample}$ at $N$ concentrations spaced $\Delta c$ apart starting at $c_1$ would require a total mass of

$$\begin{aligned} m_{tot} &= V_{sample}\left[c_1 N + \Delta c \frac{(N-1)\,N}{2}\right] \\ &= V_{sample} c_1 N \left[1 + \frac{\Delta c}{c_1}\frac{(N-1)}{2}\right]. \end{aligned} \tag{1}$$

For $V_{sample} = 0.1$ mL, $c_1 = c_{Cond} \approx 300$ mg/ml, $\Delta c = 10$ mg/ml, and $N = 3$ measurements, $m_{tot} = 93$ mg. For comparison, the yield of FUS-GFP following a typical purification is $\sim 2$ mg per liter of insect cell culture (24 aliquots $\times$ 60 $\mu$M $\times$ 20 $\mu$L $\times$ 80 kDa = 2.3 mg). While perhaps not impossible, it is highly impractical and expensive to work with on the order of $93/2.3 = 40$ L of culture. Also, typical columns are not designed for such a high material load, and purification would have to proceed in batches in series, introducing potential batch effects. The low yield and batch effects introduced by serial purification make preparation of protein samples at the high concentrations and large volumes required for bulk measurements of refractive index expensive and highly impractical in most cases.

These issues have been overcome in some limited cases. Specifically, improved yield through overexpression in bacteria and purification under denaturing conditions have enabled isolation of intrinsically disordered protein regions (IDRs) at or near the 100-mg-scale [1, 2, 3]. However, these approaches are not generically applicable to full-length proteins, particularly those containing folded domains or post-translational modifications (PTMs). Bacteria lack the machinery required for adding eukaryotic PTMs as well as the chaperones required for proper folding of many larger eukaryotic proteins. Bacterial expression systems are thus not appropriate for proteins with PTMs or that require chaperones. Further, recovery of native-state protein conformations following denaturation during purification is far from guaranteed. Following removal of chemical denaturants, proteins containing folded domains are prone to form aggregates which may not be readily reversible. Though it appears to be commonly assumed that IDRs recover a native conformational ensemble upon denaturant removal, we are not aware of a study where this equivalence has been demonstrated. The use of denaturing conditions during purification thus requires significant structural characterization and potentially extensive trial-and-error in order to attain pure protein with a native conformational ensemble, particularly when a folded domain is present. Given these limitations, measuring solution refractive index with bulk techniques at concentrations of full-length eukaryotic proteins above $c_{Cond}$ is generally not practical.

##### 1.2 Measurements below $c_{Dil}$

The primary issue with measuring $\frac{dn}{dc}$ from measurements of the refractive index $n(c)$ with $c \in [0, c_{Dil}]$ is that $c_{Dil}$ is often so low that $n(c_{Dil})$ may not be measurably different from $n(0)$. As an example, $c_{Dil}$ for PGL3-mEGFP is below 500 nM [4], corresponding to a mass concentration of $\sim 0.05$ mg/mL. Using the sequence-based prediction [5] of $\frac{dn}{dc} = 0.0001886$ mL/mg for this protein at $\lambda = 589$ nm and $T = 25°$C, the maximal increase in solution refractive index over this range is $n(c_{Dil}) - n(0) = \frac{dn}{dc} c_{Dil} \approx 9 \times 10^{-6}$. This is smaller than the sensitivity of our research-grade digital refractometer, which (only) measures out to five digits.

Given the impossibility of measuring $\frac{dn}{dc}$ at protein concentrations within the miscibility gap, the exceptional sensitivity required to measure it at low concentrations, and the impracticality of measuring it at high concentrations, we find the sequence-based estimates adequate for current purposes.

#### 2 Note 2: Linear approximation to refractive index of mixtures

There is an extensive literature regarding appropriate "mixing rules" for calculating the refractive index of mixtures [6, 7, 8, 9], which in general depend non-linearly on concentration. Recognizing this, we make no claim regarding the universality of the approximation used in this paper, namely that the refractive index of a mixture is given by a linear sum of contributions from the non-solvent components. Rather, we emphasize that the refractive index of aqueous solutions of biomolecules often shows a linear concentration dependence over a wide range of concentrations [10, 11, 12, 13].

As checks for the validity of the approximation in the current work, we demonstrate linearity explicitly for the protein BSA, the linear polymer PEG, the branched polymer Dextran, and the nucleic acid polyA RNA (Fig. 2i, Fig. S3). As a check on the linear sum approximation for multi-component mixtures, we compared the refractive index estimated from the linear sum approximation to measurements of homogeneous PEG/Dextran mixtures and found only small deviations between the two (Fig. S5). These data thus support the use of the linear sum approximation (Main Text Eq. 3) for the systems studied in this work.

### 3 Note 3: Measurement of $c_{Dil}$

The final parameter in Eq (2) from the Main Text is $c_{Dil}$. If sufficiently large, $c_{Dil}$ can be determined from the refractive index of the dilute phase, $n_{Dil}$. Since the dilute phase accounts for $> 99$ % of the sample volume in typical biomolecular condensate reconstitutions, it is typically possible to produce (e.g. via centrifugation) the ∼100 $\mu$L of pure dilute phase required to measure $n_{Dil}$ with bulk techniques like digital refractometry. The concentration threshold for $c_{Dil}$ measurements at 2-digits of precision with a typical digital refractometer is about 0.5 mg/ml (see also section 1.2). For $c_{Dil} < c_{thresh} \approx 0.5$ mg/ml, standard analytic chemistry techniques (e.g. UV-VIS spectroscopy, mass spectrometry, or Western blotting) may be required to determine $c_{Dil}$. However, as we find $c_{Cond}$ is typically $\gtrsim 100$ mg/ml, the contribution of $c_{Dil}$ to $c_{Cond}$ is typically negligible, particularly when $c_{Dil}$ is too low to detect by refractometery. The contribution of $c_{Dil}$ in Eq. (2) from the Main Text may therefore often be neglected.

### 4 Note 4: Requirement that tie-lines are straight

The goal of this section is to demonstrate that thermodynamic tie-lines in multi-component systems are necessarily linear relationships between component concentrations, and therefore are represented in phase diagrams by straight lines (when plotted on linear axes). We proceed by introducing necessary definitions, and then showing that this linear relationship follows directly from mass conservation and incompressibility.

Consider a closed system of total volume $V^{tot}$ filled with molecules of N distinct chemical species. Let $c_i$, $M_{W,i}$, and $\overline{\nu}_i$ be the molar concentration (mole/liter), molar mass (grams/mole), and partial specific volume (liter/gram) of component $i$. The average system composition is uniquely specified by $\overline{\phi} = \{\overline{\phi}_1, ..., \overline{\phi}_i, ..., \overline{\phi}_N\}$, where $\overline{\phi}_i = c_i M_{W,i} \overline{\nu}_i$ is the average volume fraction of species $i$.

Without loss of generality, we assume that the chemical species are not all mutually miscible at all proportions. That is, we assume that there is a contiguous region in the N-dimensional composition-space spanned by $\overline{\phi}$ in which the system demixes into two or more phases. For simplicity, we restrict ourselves here to the case where demixing results in exactly two phases. In this case, the region of the miscibility gap is bounded by an (N-1)-dimensional manifold called the binodal. Since this manifold is either closed or bounded by a subset of the $(\phi_i, \phi_j)$-planes defining the outer edges of composition space, the notions of “inside” and “outside” the binodal are unambiguously defined. For $\overline{\phi}$ values outside the binodal, the system exists in equilibrium as a single homogeneous phase whose composition is given by $\overline{\phi}$. For $\overline{\phi}$ inside the binodal, however, the system consists, in equilibrium, of two co-existing phases labeled I and II, the compositions of which are given by points $\phi^I$ and $\phi^{II}$ on the binodal. The collection of points $\overline{\phi}$ in the immiscible region which map to the same two points $\phi^I$ and $\phi^{II}$ on the binodal form a contiguous curve called a tie-line.

To show that there is a mathematical relationship between the points $\overline{\phi}$, $\phi^I$, and $\phi^{II}$ on a tie-line, we start by noting that mass concervation requires that the total mass of component $i$ in the system equal the sum of the mass of $i$ in each of the phases

$$M_i^{tot} = M_i^I + M_i^{II} \,. \tag{2}$$

Since the mass of component $i$ in a volume $V$ is given by $M_i = c_i M_{W,i} V = \phi_i V / \overline{\nu}_i$, (2) becomes

$$\overline{\phi}_i V^{tot} = \phi_i^I V^I + \phi_i^{II} V^{II} \,, \tag{3}$$

where $V^I$ and $V^{II}$ are the volumes of the two phases. Eq. (3) represents a relationship between the amount of a single component in the entire system and the amounts in each phase.

Before we can demonstrate a relationship between these values for multiple components simultaneously, it is useful to first simplify Eq. (3) by removing the dependence on $V^{II}$. To this end, we assume that the system is incompressible, such that

$$V^{tot} = V^I + V^{II} \,. \tag{4}$$

With this constraint, Eq. (3) can be rewritten as

$$V^I = \frac{\phi_i^{II} - \overline{\phi}_i}{\phi_i^{II} - \phi_i^I} V^{tot} \,, \tag{5}$$

a statement of the well-known lever-rule [14] that gives the volume of a phase in terms of the amounts of a component in each phase as well as the system as a whole.

To obtain a relationship between compositions of two different components, we note that applications of Eq. (5) with components $i \neq j$ must give the same phase volume

$$\frac{\phi_i^{II} - \overline{\phi}_i}{\phi_i^{II} - \phi_i^{I}} V^{tot} = V^{I} = \frac{\phi_j^{II} - \overline{\phi}_j}{\phi_j^{II} - \phi_j^{I}} V^{tot} . \tag{6}$$

Eq. (6) can be rearranged to obtain an expression for the average volume fraction of one component in the immiscible region in terms of the average volume fraction of a second component

$$\overline{\phi}_j = \left[ \frac{\phi_j^{II} - \phi_j^{I}}{\phi_i^{II} - \phi_i^{I}} \right] \overline{\phi}_i + \left[ \phi_j^{II} - \left( \frac{\phi_j^{II} - \phi_j^{I}}{\phi_i^{II} - \phi_i^{I}} \right) \phi_i^{II} \right] . \tag{7}$$

Importantly, Eq. (7) provides a linear relationship between $\overline{\phi}_j$ and $\overline{\phi}_i$, written here in the form of a straight line in the $(\phi_i, \phi_j)$-plane with slope $m = \frac{\phi_j^{II} - \phi_j^{I}}{\phi_i^{II} - \phi_i^{I}}$ and y-intercept $b = \phi_j^{II} - \left( \frac{\phi_j^{II} - \phi_j^{I}}{\phi_i^{II} - \phi_i^{I}} \right) \phi_i^{II}$. Note that both the slope and y-intercept depend only on the coordinates of the points on the binodal, and are thus constant for all $\overline{\phi}$ on the same tie-line. This indicates that the projection of the tie-line into the $(\phi_i, \phi_j)$-plane is linear. Since Eq. (7) is linear for any choice of two distinct components $i$ and $j$, it follows that the tie-line must appear linear following projection into any plane spanned by a pair of distinct components. Furthermore, since the only curve for which every projection is linear is itself linear, we conclude that the tie-line must be a linear (i.e. a straight line) function embedded in the N-dimensional space spanned by $\phi$, as claimed.

### 5 Note 5: Derivation of condensate composition for multi-component systems

#### 5.1 Overview

In this supplementary note, we derive Eq. 4 in the main text. In the first section, we explicitly consider a ternary mixture composed of one solvent and two solutes. We construct a linear system of constraint equations, invert them, and present the solution. In the second section, we repeat this procedure in the context of a quaternary system (1 solvent + 3 solutes). We provide expressions in forms that readily generalize to systems with more components, and comment on selection of tie-line projections and requirements for the matrix representing the linear system to be invertible. In the third section, we briefly present the the calculation for a mixture with an arbitrary number of components (1 solvent + $N$ solutes). In the fourth section, we outline the procedure used to calculate the uncertainty in the condensed-phase concentrations obtained with this method.

#### 5.2 Ternary systems

In this section, we study the regime of two-phase coexistence of a ternary mixture at constant temperature. Specifically, we derive an explicit expression for the composition of one phase in terms of the composition of the other, the system average composition, and the refractive index increments of the solutes.

Consider a closed system of total volume $V^{tot}$ containing three distinct chemical species, a solvent and $N = 2$ solutes, indexed by $i \in \{0, 1, 2\}$. In the context of the experiments of Fig. 5 in the main text, these species would be an effective solvent representing the aqueous buffer, the protein FUS-GFP and polyA RNA. Similar to section 4 above, let $M_{W,i}$ and $\overline{\nu}_i$ be the molar mass (grams/mole), and partial specific volume (liter/gram) of species $i$, but with $c_i$ now the average mass concentration (grams/liter) of species $i$. The average system composition is uniquely specified by $\overline{\mathbf{c}} = \{\overline{c}_0, \overline{c}_1, \overline{c}_2\}$.

We start by assuming the existence of a two-phase coexistence region in the composition space spanned by $\overline{\mathbf{c}}$, i.e. that the three components are not mutually miscible in all proportions. Though the Gibbs Phase Rule permits the existence of a three-phase coexistence region, we restrict ourselves in the following to consider $\overline{\mathbf{c}}$ only in the two-phase region. We label the phases with Roman numerals, indexed by $\alpha$. Without loss of generality, we assume that the concentration of component 1 is higher in phase II than in phase I, i.e. $c_1^I < \overline{c}_1 < c_1^{II}$. Relative to component 1, phase I thus corresponds to the dilute phase, while phase II corresponds to the condensed phase.

Given an average composition $\overline{\mathbf{c}} = \mathbf{c^B}$ in the two-phase regime and the corresponding dilute-phase composition $\mathbf{c^I} = \mathbf{c^A}$, our goal is to calculate the composition of the coexisting condensed phase, $\mathbf{c^{II}}$. To uniquely determine the concentrations of all three components at $\mathbf{c^{II}} \equiv \{c_0^{II}, c_1^{II}, c_2^{II}\}$, we require three linearly independent equations relating these concentrations to other known values. The first equation come from assuming both phases to be incompressible, such that Eq. 4 holds. Specifically, incompressibility implies that the sum of component volume fractions in each phase $\alpha$ sum to unity,

$$\sum_i \phi_i^\alpha = \sum_i c_i^\alpha \overline{\nu}_i = 1. \tag{8}$$

From this relationship, the concentration of component $i$ in phase $\alpha$ can be calculated immediately if the other two are known:

$$c_i^\alpha = \overline{\nu}_i^{-1}(1 - \sum_{j \neq i} c_j^\alpha \overline{\nu}_j). \tag{9}$$

In the following, we assume component 0 to be the (effective) solvent, and use Eq. 9 to determine $c_0$ in each phase. The problem of calculating $\mathbf{c^{II}}$ in a ternary system is thus reduced to determining $c_1^{II}$ and $c_2^{II}$. This is the situation described graphically in Fig. 5a of the main text in the context of a phase diagram in which species 1 and 2 are both enriched in phase $II$. We note, however, that the approach does not require that species 2 also be enriched in phase $II$.

As indicated above, we require two additional constraints to proceed. Physically, these constraints come from measurements of the refractive index difference between the two co-existing phases by QPI, and the thermodynamic tie-line connecting $\mathbf{c^I}$ to $\mathbf{c^{II}}$. Approximating the refractive index difference as a linear combination of contributions from concentration differences in each non-solvent component, we can express it as

$$\Delta n = \frac{dn}{dc_1}(c_1^{II} - c_1^I) + \frac{dn}{dc_2}(c_2^{II} - c_2^I), \tag{10}$$

where $\frac{dn}{dc_i}$ is the refractive index increment of component $i$. By rearranging Eq. 10 slightly and replacing $(c_1^{II}, c_2^{II})$ with the generic point $(c_1, c_2)$, we arrive at the form of a straight line in the $(c_1, c_2)$-plane:

$$\begin{aligned} c_2 &= -\left(\frac{\frac{dn}{dc_1}}{\frac{dn}{dc_2}}\right) c_1 + \left(\frac{1}{\frac{dn}{dc_2}}\left(\Delta n + \frac{dn}{dc_1} c_1^I\right) + c_2^I\right) \\ &= m_{IRL} c_1 + b_{IRL}. \end{aligned} \tag{11}$$

Physically, Eq. 11 represents a line of constant refractive index difference (isorefractive line) in the $(c_1, c_2)$-plane. This can be seen by noting that each point $(c_1, c_2)$ satisfying Eq. 11 specifies a potential condensed-phase composition for which the refractive index difference between it and the dilute phase $(c_1^I, c_2^I)$ is consistent with the measured value $\Delta n$.

For the final constraint, we construct the thermodynamic tie-line connecting $\mathbf{c^I}$ to $\mathbf{c^{II}}$ in the $(c_1, c_2)$-plane. The tie-line slope can be calculated directly from the coordinates of $\mathbf{c^I}$ and $\overline{\mathbf{c}}$ as

$$m_{TL} = \frac{\overline{c}_2 - c_2^I}{\overline{c}_1 - c_1^I}, \tag{12}$$

while the $y$-intercept is given by

$$b_{TL} = \overline{c}_2 - m_{TL}\overline{c}_1. \tag{13}$$

The equation for the tie-line in the $(c_1, c_2)$-plane may therefore be written as

$$c_2 = m_{TL} c_1 + (\overline{c}_2 - m_{TL}\overline{c}_1). \tag{14}$$

Since the point $(c_1^{II}, c_2^{II})$ must lie on the tie-line by definition, our final constraint is thus

$$c_2^{II} = m_{TL} c_1^{II} + (\overline{c}_2 - m_{TL}\overline{c}_1). \tag{15}$$

Taken together, equations 10 and 15 suffice to fully specify a system of linear equations in the unknowns $(c_1^{II}, c_2^{II})$. We now cast this in matrix form as

$$\begin{bmatrix} \frac{dn}{dc_1} & \frac{dn}{dc_2} \\ m_{TL} & -1 \end{bmatrix} \begin{bmatrix} c_1^{II} \\ c_2^{II} \end{bmatrix} = \begin{bmatrix} \Delta n + \frac{dn}{dc_1} c_1^I + \frac{dn}{dc_2} c_2^I \\ m_{TL}\bar{c}_1 - \bar{c}_2 \end{bmatrix} \tag{16}$$

$$M\mathbf{c^{II}} = \mathbf{x}. \tag{17}$$

With the identification $(c_1^{II}, c_2^{II}) \to (p^{II}, r^{II})$, Eq. 16 is equivalent to that presented in Fig. 5b in the main text. To assess invertability of the matrix $M$, we next calculate its determinant,

$$det(M) = -\frac{dn}{dc_1} - m_{TL}\frac{dn}{dc_2}. \tag{18}$$

Noting that the refractive index increments for most biomolecules in water are positive constants, the matrix $M$ is singular (non-invertible) only if $m_{TL} = -\frac{dn}{dc_1}/\frac{dn}{dc_2} = m_{IRL}$. The indeterminacy stems from the fact that, in this special case, the isorefractive line and the tie-line would be coincident; Equations 10 and 15 would describe the same line and thus not provide linearly independent information. Such a special case would correspond strictly to segregative phase separation (species 1 and 2 are enriched in opposite phases), and then only in regions of the phase diagram that happen to have the misfortune of containing tie-lines with the same slope as the isorefractive line. In practice, we expect this situation to arise only rarely. In cases where matrix singularity is an issue, a change in system temperature may be sufficient to shift $det(M)$ away from the singularity by altering the tie-line slopes relative to the isorefractive line.

For the typical case of non-singular $M$, the system inverse is given by

$$M^{-1} = \frac{1}{det(M)} \begin{bmatrix} -1 & -\frac{dn}{dc_2} \\ -m_{TL} & \frac{dn}{dc_1} \end{bmatrix}. \tag{19}$$

With this, the condensed-phase composition is given by

$$\mathbf{c^{II}} = M^{-1}\mathbf{x}. \tag{20}$$

After some algebra, and noting that

$$c_2^I - \bar{c}_2 + \bar{c}_1 m_{TL} = c_1^I m_{TL} \tag{21}$$

$$c_1^I - \bar{c}_1 + \frac{\bar{c}_2}{m_{TL}} = \frac{c_2^I}{m_{TL}}, \tag{22}$$

Eq. 20 simplifies to

$$\begin{bmatrix} c_1^{II} \\ c_2^{II} \end{bmatrix} = \begin{bmatrix} \frac{\Delta n}{-det(M)} + c_1^I \\ \frac{\Delta n m_{TL}}{-det(M)} + c_2^I \end{bmatrix}. \tag{23}$$

Equation 23 was used to compute the condensed-phase concentrations of FUS-GFP and polyA RNA reported in Fig. 5. With $c_1^{II}$ and $c_2^{II}$ determined by Eq. 24, the solvent concentration $c_0^{II}$ can be calculated from Eq. 9. The condensed-phase composition is thus fully specified.

We note that, from comparison to Eq. 19, the first column of $M^{-1}$ appears on the right-hand side of Eq. 23. We can thus rewrite Eq. 23 in for the individual components $c_i^{II}$ as

$$c_i^{II} = \Delta n M_{i1}^{-1} + c_i^I, \tag{24}$$

where $M_{i1}^{-1}$ is the $i$th matrix element of the first column of $M^{-1}$. Equation 24 is identical to Eq. 4 given in the main text, and, as we'll see below, is the form of the general solution for the condensed-phase composition of the $N$ solutes in an $N+1$-component mixture.

#### 5.3 Quaternary systems

In a quaternary system (4 species), we have 1 solvent and $N = 3$ solutes indexed by $i \in \{0, 1, 2, 3\}$. As for the ternary system, we restrict ourselves to the region of the space spanned by $\bar{\mathbf{c}}$ corresponding to two-phase coexistence. We also assume, without loss of generality, that species 1 is enriched in phase $II$. We'll again use Eq. 9 to determine $c_0^{II}$, so we need three constraints to determine the condensed-phase concentrations of the other three components. The first constraint is the quaternary analog of Eq. 10 for the refractive index difference between the coexisting phases,

$$\Delta n = \frac{dn}{dc_1}(c_1^{II} - c_1^I) + \frac{dn}{dc_2}(c_2^{II} - c_2^I) + \frac{dn}{dc_3}(c_3^{II} - c_3^I), \tag{25}$$

which now corresponds to an isorefractive plane in the space spanned by $\langle c_1, c_2, c_3 \rangle$.

The other two constraints come from projections of the thermodynamic tie-line into 2-dimensional subspaces. In a quaternary system with $N = 3$ solutes, there are $\binom{N}{2} = 3$ such projections possible: the $(c_1, c_2)$-plane, the $(c_1, c_3)$-plane, and the $(c_2, c_3)$-plane. However, only $N - 1 = 2$ of these are linearly independent. We could therefore choose any two. We select here the projections into the $(c_1, c_2)$-plane and the $(c_1, c_3)$-plane, as the assumption that component 1 is enriched in phase $II$ guarantees that the slopes of the projections are finite.

Following the procedure for ternary systems above, the slope for the tie-line projected into the $(c_i, c_j)$-plane is

$$m_{ij} = \frac{\bar{c}_j - c_j^I}{\bar{c}_i - c_i^I}. \tag{26}$$

Along with the $y$-intercept

$$b_{ij} = \bar{c}_j - m_{ij}\bar{c}_i, \tag{27}$$

the equation for the projection of the thermodynamic tie-line into the $(c_i, c_j)$-plane is given by

$$c_j = m_{ij}c_i + b_{ij}. \tag{28}$$

We next combine the constraints from the isorefractive plane (Eq. 25) and the tie-line projections into the $(c_1, c_2)$- and $(c_1, c_3)$-planes into a matrix equation of the form $M\mathbf{c^{II}} = \mathbf{x}$ as

$$\begin{bmatrix} \frac{dn}{dc_1} & \frac{dn}{dc_2} & \frac{dn}{dc_3} \\ m_{12} & -1 & 0 \\ m_{13} & 0 & -1 \end{bmatrix} \begin{bmatrix} c_1^{II} \\ c_2^{II} \\ c_3^{II} \end{bmatrix} = \begin{bmatrix} \Delta n + \sum\limits_{i \neq 0} \frac{dn}{dc_i} c_i^I \\ -b_{12} \\ -b_{13} \end{bmatrix}. \tag{29}$$

To assess whether the system can be solved, we again calculate the determinant of $M$, which is given by

$$det(M) = \frac{dn}{dc_1} + \frac{dn}{dc_2}m_{12} + \frac{dn}{dc_3}m_{13} = \frac{dn}{dc_1} + \sum_{i>1} \frac{dn}{dc_i}m_{1i}. \tag{30}$$

As always, $det(M)$ must be finite and non-zero for $M$ to be invertible. Setting Eq. 30 equal to zero yields a condition on $m_{13}$ for which $M$ is singular and non-invertible, namely

$$m_{13} = -\left( \frac{\frac{dn}{dc_1}}{\frac{dn}{dc_3}} + \frac{\frac{dn}{dc_2}}{\frac{dn}{dc_3}} m_{12} \right). \tag{31}$$

Similar to the ternary case, this condition corresponds to the tie-line lying in the isorefractive plane defined by Eq. 25. Since the $m_{ij}$ are generally independent of each other and also physically independent of the refractive index increments, there is no physical reason to expect this to occur often. Therefore, so long as the tie-line slopes do not by chance happen to satisfy the relationship given in Eq. 31, $M$ is invertible.

We briefly mention a few specific and potentially common cases. First, we note that if the $m_{ij}$ are both positive, $det(M)$ is positive and the system is always invertible. This corresponds to components 2 and 3 also being enriched in phase $II$. Second, if $m_{ij} = 0$ for one of the components j, then the concentration of that component in each phase is given by $c_j^{II} = c_j^I = \bar{c}_j$ (i.e. its partition coefficient is 1) and the situation simplifies to that of the ternary case discussed above. Third, if $m_{ij} = 0$ for both components 2 and 3, then the situation reduces to an effective binary system and can be solved from Eq. 25 alone.

Assuming the tie-line is such that $M$ is non-singular, its inverse is given by

$$M^{-1} = \frac{1}{det(M)} \begin{bmatrix} 1 & \frac{dn}{dc_2} & \frac{dn}{dc_3} \\ m_{12} & -\frac{dn}{dc_1} - \frac{dn}{dc_3} m_{13} & \frac{dn}{dc_3} m_{12} \\ m_{13} & \frac{dc}{dc_2} m_{13} & -\frac{dn}{dc_1} - \frac{dn}{dc_2} m_{12} \end{bmatrix} . \tag{32}$$

The condensed-phase composition can thus be computed from $\mathbf{c^{II}} = M^{-1}\mathbf{x}$. After some algebra, this simplifies to

$$\begin{bmatrix} c_1^{II} \\ c_2^{II} \\ c_3^{II} \end{bmatrix} = \begin{bmatrix} \frac{\Delta n}{det(M)} + c_1^I \\ \frac{\Delta n m_{12}}{det(M)} + c_2^I \\ \frac{\Delta n m_{13}}{det(M)} + c_3^I \end{bmatrix} . \tag{33}$$

As in the Ternary case, comparison of Eq. 33 to Eq. 32 shows that the first column of $M^{-1}$ again shows up in the solution. We thus find that

$$c_i^{II} = \Delta n M_{i1}^{-1} + c_i^I \tag{34}$$

for Quaternary systems as well. Physically, the matrix elements of the first column of $M^{-1}$ denote the contribution of each component to the refractive index difference between the two phases.

#### 5.4 General solution for (N+1)-component systems

The procedure for a solution with 1 solvent and $N$ solute species follows directly from the results for the quaternary mixtures of the previous section. The refractive index difference between phases

$$\Delta n = \sum_{i \neq 0} \frac{dn}{dc_i} \left( c_i^{II} - c_i^{II} \right) \tag{35}$$

represents a hyperplane in the $N$-dimensional space spanned by the solute concentrations. Generating tie-line constraints from projections of the thermodynamic tie-line into the $(c_1, c_j)$-plane for all $N-1$ choices of $j$, the constraints can be written together in a matrix expression of the form $M\mathbf{c^{II}} = \mathbf{x}$ as

$$\begin{bmatrix} \frac{dn}{dc_1} & \frac{dn}{dc_2} & \frac{dn}{dc_3} & \dots & \frac{dn}{dc_i} & \dots & \frac{dn}{dc_N} \\ m_{12} & -1 & 0 & \dots & 0 & \dots & 0 \\ m_{13} & 0 & -1 & \dots & 0 & \dots & 0 \\ \vdots & \vdots & \vdots & \ddots & \vdots & \vdots & \vdots \\ m_{1i} & 0 & 0 & \dots & -1 & \dots & 0 \\ \vdots & \vdots & \vdots & \vdots & \vdots & \ddots & \vdots \\ m_{1N} & 0 & 0 & \dots & 0 & \dots & -1 \end{bmatrix} \begin{bmatrix} c_1^{II} \\ c_2^{II} \\ c_3^{II} \\ \vdots \\ c_i^{II} \\ \vdots \\ c_N^{II} \end{bmatrix} = \begin{bmatrix} \Delta n + \sum_{i \neq 0} \frac{dn}{dc_i} c_i^I \\ -b_{12} \\ -b_{13} \\ \vdots \\ -b_{1i} \\ \vdots \\ -b_{1N} \end{bmatrix} . \tag{36}$$

The determinant of $M$ is given again by

$$det(M) = \frac{dn}{dc_1} + \sum_{i>1} \frac{dn}{dc_i} m_{1i}, \tag{37}$$

and is zero if and only if the thermodynamic tie-line lies in the isorefractive hyperplane. As the number of solutes $N$ increases, the likelihood of this happening by chance becomes progressively smaller.

The inverse of $M$ in the general case is given by

$$M^{-1} = \frac{1}{det(M)} \times$$

$$\begin{bmatrix} 1 & \frac{dn}{dc_2} & \frac{dn}{dc_3} & \cdots & \frac{dn}{dc_i} & \cdots & \frac{dn}{dc_N} \\ m_{12} & -\frac{dn}{dc_1} - \sum\limits_{1<k\neq 2} \frac{dn}{dc_k} m_{1k} & \frac{dn}{dc_3} m_{12} & \cdots & \frac{dn}{dc_i} m_{12} & \cdots & \frac{dn}{dc_N} m_{12} \\ m_{13} & \frac{dn}{dc_2} m_{13} & -\frac{dn}{dc_1} - \sum\limits_{1<k\neq 3} \frac{dn}{dc_k} m_{1k} & \cdots & \frac{dn}{dc_i} m_{13} & \cdots & \frac{dn}{dc_N} m_{13} \\ \vdots & \vdots & \vdots & \ddots & \vdots & \vdots & \vdots \\ m_{1i} & \frac{dn}{dc_2} m_{1i} & \frac{dn}{dc_3} m_{1i} & \cdots & -\frac{dn}{dc_1} - \sum\limits_{1<k\neq i} \frac{dn}{dc_k} m_{1k} & \cdots & \frac{dn}{dc_N} m_{1i} \\ \vdots & \vdots & \vdots & \vdots & \vdots & \ddots & \vdots \\ m_{1N} & \frac{dn}{dc_2} m_{1N} & \frac{dn}{dc_3} m_{1N} & \cdots & \frac{dn}{dc_i} m_{1N} & \cdots & -\frac{dn}{dc_1} - \sum\limits_{1<k\neq N} \frac{dn}{dc_k} m_{1k} \end{bmatrix}. \tag{38}$$

Using this expression for $M^{-1}$ in $\mathbf{c^{II}} = M^{-1}\mathbf{x}$, the condensed-phase composition for the general case simplifies (after much algebra) to

$$\begin{bmatrix} c_1^{II} \\ c_2^{II} \\ c_3^{II} \\ \vdots \\ c_i^{II} \\ \vdots \\ c_N^{II} \end{bmatrix} = \begin{bmatrix} \frac{\Delta n}{det(M)} + c_1^I \\ \frac{\Delta n m_{12}}{det(M)} + c_2^I \\ \frac{\Delta n m_{13}}{det(M)} + c_3^I \\ \vdots \\ \frac{\Delta n m_{1i}}{det(M)} + c_i^I \\ \vdots \\ \frac{\Delta n m_{1N}}{det(M)} + c_N^I \end{bmatrix}, \tag{39}$$

or, more compactly, as the now familiar expression

$$c_i^{II} = \Delta n M_{i1}^{-1} + c_i^I. \tag{40}$$

As the number of solute components $N$ is arbitrary throughout this section, Eq. 40 may be used to calculate the species concentrations in phase $II$ for mixtures with an arbitrary number of components, as claimed in the main text.

#### 5.5 Error propagation with Jacobians

In this section we document how we calculated the uncertainty on the condensed-phase concentrations we report for the ternary FUS/RNA system in Fig. 5. The condensed-phase concentrations are calculated from Eq. 40. For each component $i$, Eq. 40 prescribes a function of seven variables $f_i\left(\Delta n, \frac{dn}{dc_1}, \frac{dn}{dc_2}, \bar{c}_1, \bar{c}_2, c_1^I, c_2^I, \right)$. Measurement or estimation of each variable $x$ comes with an associated uncertainty $\sigma_x$. We estimate the variance of $f_i$ as

$$(\delta f_i)^2 = J_i \Sigma^x J_i^T, \tag{41}$$

where $J_i$ is the Jacobian of $f_i$. $J_i$ is a row vector given by

$$J_i = \begin{bmatrix} \frac{\partial f_i}{\partial \Delta n} & \frac{\partial f_i}{\partial \left(\frac{dn}{dc_1}\right)} & \frac{\partial f_i}{\partial \left(\frac{dn}{dc_2}\right)} & \frac{\partial f_i}{\partial \bar{c}_1} & \frac{\partial f_i}{\partial \bar{c}_2} & \frac{\partial f_i}{\partial c_1^I} & \frac{\partial f_i}{\partial c_2^I} \end{bmatrix}, \tag{42}$$

and $J^T$ is its transpose. $\Sigma^x$ is a square matrix containing the cross-correlations $\rho_{jk}$ between the uncertainties of the variables $x_j$ and $x_k$ weighted by the product of their respective uncertainties $\sigma_j \sigma_k$. The matrix elements of $\Sigma^x$ are thus given by

$$\Sigma_{jk}^x = \rho_{jk} \sigma_j \sigma_k. \tag{43}$$

Final error bars on $f_i$ would represent the standard deviation $\delta f_i$.

Since each variable $x_j$ was determined independently for the FUS/RNA measurements in Fig 5, we used a diagonal cross-correlation matrix where $\rho_{jj} = 1$ and $\rho_{jk} = 0$ for $j \neq k$. We next list how the uncertainties $\sigma_x$ in each variable $x$ were determined. For $\sigma_{\Delta n}$, we used the standard deviation of the $\Delta n$ measurements from the corresponding condition. For $\sigma_{dn/dc_2}$, we used the 95 %-confidence interval from the linear fit in Fig. S4. Since $\frac{dn}{dc_1}$ was estimated from sequence rather than measured directly, we estimated $\sigma_{dn/dc_1}$ as $\left(\frac{dn/dc_1}{dn/dc_2}\right)\sigma_{dn/dc_2}$. $\sigma_{\bar{c}_1}$ and $\sigma_{\bar{c}_1}$ represent the standard deviations of repeat spectrophotometric measurements of the stock concentrations, rescaled by the ratio $\bar{c}_i/c_i^{stock}$. Finally, $\sigma_{c_1^I}$ and $\sigma_{c_1^I}$ are proportional to standard deviations from repeat measurements of the dilute-phase UV-Vis spectra.

For calculating the variance of the condensed-phase protein concentration $\left(\delta c_1^{II}\right)^2$ from Eq. 41, we used

$$\begin{aligned} f_1 &= \frac{\Delta n}{-det(M)} + c_1^I \\ &= \frac{\Delta n}{\frac{dn}{dc_1} + \frac{dn}{dc_2}\frac{\bar{c}_2 - c_2^I}{\bar{c}_1 - c_1^I}} + c_1^I, \end{aligned} \tag{44}$$

for which the Jacobian reads

$$J_1 = -\frac{\Delta n}{\left(det(M)\right)^2}\begin{bmatrix} \frac{det(M)}{\Delta n} & 1 & m_{TL} & \frac{m_{TL}\frac{dn}{dc_2}}{\left(\bar{c}_1 - c_1^I\right)} & -\frac{\frac{dn}{dc_2}}{\left(\bar{c}_1 - c_1^I\right)} & \frac{m_{TL}\frac{dn}{dc_2}}{\left(\bar{c}_1 - c_1^I\right)} - 1 & -\frac{\frac{dn}{dc_2}}{\left(\bar{c}_1 - c_1^I\right)} \end{bmatrix}, \tag{45}$$

with

$$det(M) = -\frac{dn}{dc_1} - \frac{dn}{dc_2}\frac{\bar{c}_2 - c_2^I}{\bar{c}_1 - c_1^I}, \tag{46}$$

and

$$m_{TL} = \frac{\bar{c}_2 - c_2^I}{\bar{c}_1 - c_1^I}. \tag{47}$$

For calculating the variance of the condensed-phase RNA concentration $\left(\delta c_2^{II}\right)^2$ from Eq. 41, we used

$$\begin{aligned} f_2 &= \frac{\Delta n m_{TL}}{-det(M)} + c_2^I \\ &= \frac{\Delta n\left(\frac{\bar{c}_2 - c_2^I}{\bar{c}_1 - c_1^I}\right)}{\frac{dn}{dc_1} + \frac{dn}{dc_2}\frac{\bar{c}_2 - c_2^I}{\bar{c}_1 - c_1^I}} + c_2^I \\ &= \frac{\Delta n}{\frac{dn}{dc_1}\left(\frac{\bar{c}_1 - c_1^I}{\bar{c}_2 - c_2^I}\right) + \frac{dn}{dc_2}} + c_2^I \\ &= \frac{\Delta n}{\Omega} + c_2^I \end{aligned}, \tag{48}$$

with

$$\Omega \equiv \frac{dn}{dc_1}\left(\frac{\bar{c}_1 - c_1^I}{\bar{c}_2 - c_2^I}\right) + \frac{dn}{dc_2}. \tag{49}$$

In this case, the Jacobian reads

$$J_2 = \frac{\Delta n}{\Omega^2}\begin{bmatrix} \frac{\Omega}{\Delta n} & -\frac{1}{m_{TL}} & -1 & -\frac{\frac{dn}{dc_1}}{\left(\bar{c}_2 - c_2^I\right)} & \frac{\frac{dn}{dc_1}}{m_{TL}\left(\bar{c}_2 - c_2^I\right)} & \frac{\frac{dn}{dc_1}}{\left(\bar{c}_2 - c_2^I\right)} & 1 - \frac{\frac{dn}{dc_1}}{m_{TL}\left(\bar{c}_2 - c_2^I\right)} \end{bmatrix}. \tag{50}$$

The final error bars on the condensed-phase protein concentrations $c_1^{II}$ in Fig. 5F represent the standard deviation $\delta c_1^{II}$ calculated according to

$$\delta c_1^{II} = \sqrt{J_1 \Sigma^x J_1^T}. \tag{51}$$

Similarly, the final error bars on the condensed-phase RNA concentrations $c_2^{II}$ in Fig. 5F represent the standard deviation $\delta c_2^{II}$ calculated according to

$$\delta c_2^{II} = \sqrt{J_2 \Sigma^x J_2^T}. \tag{52}$$

### 6 Bibliography

[1] Jacob P. Brady, Patrick J. Farber, Ashok Sekhar, Yi-Hsuan Lin, Rui Huang, Alaji Bah, Timothy J. Nott, Hue Sun Chan, Andrew J. Baldwin, Julie D. Forman-Kay, and Lewis E. Kay. Structural and hydrodynamic properties of an intrinsically disordered region of a germ cell-specific protein on phase separation. *Proceedings of the National Academy of Sciences*, 114(39):E8194–E8203, September 2017.

[2] Erik W. Martin, Alex S. Holehouse, Ivan Peran, Mina Farag, J. Jeremias Incicco, Anne Bremer, Christy R. Grace, Andrea Soranno, Rohit V. Pappu, and Tanja Mittag. Valence and patterning of aromatic residues determine the phase behavior of prion-like domains. *Science*, 367(6478):694–699, February 2020. Publisher: American Association for the Advancement of Science Section: Report.

[3] Anne Bremer, Mina Farag, Wade M. Borcherds, Ivan Peran, Erik W. Martin, Rohit V. Pappu, and Tanja Mittag. Deciphering how naturally occurring sequence features impact the phase behaviours of disordered prion-like domains. *Nature Chemistry*, 14(2):196–207, February 2022. Number: 2 Publisher: Nature Publishing Group.

[4] Shambaditya Saha, Christoph A. Weber, Marco Nousch, Omar Adame-Arana, Carsten Hoege, Marco Y. Hein, Erin Osborne-Nishimura, Julia Mahamid, Marcus Jahnel, Louise Jawerth, Andrej Pozniakovski, Christian R. Eckmann, Frank Jülicher, and Anthony A. Hyman. Polar Positioning of Phase-Separated Liquid Compartments in Cells Regulated by an mRNA Competition Mechanism. *Cell*, 166(6):1572–1584.e16, September 2016.

[5] Huaying Zhao, Patrick H. Brown, and Peter Schuck. On the Distribution of Protein Refractive Index Increments. *Biophysical Journal*, 100(9):2309–2317, May 2011.

[6] Wilfried Heller. Remarks on Refractive Index Mixture Rules. *The Journal of Physical Chemistry*, 69(4):1123–1129, April 1965.

[7] Thomas G. Mayerhöfer, Alicja Dabrowska, Andreas Schwaighofer, Bernhard Lendl, and Jürgen Popp. Beyond Beer's Law: Why the Index of Refraction Depends (Almost) Linearly on Concentration. *ChemPhysChem*, 21(8):707–711, 2020. _eprint: https://chemistry-europe.onlinelibrary.wiley.com/doi/pdf/10.1002/cphc.202000018.

[8] Huib J. Bakker. Reaction-field model for the dielectric response of mixtures. *The Journal of Chemical Physics*, 153(5):054503, August 2020. Publisher: American Institute of Physics.

[9] Bilin Zhuang, Gabriele Ramanauskaite, Zhao Yuan Koa, and Zhen-Gang Wang. Like dissolves like: A first-principles theory for predicting liquid miscibility and mixture dielectric constant. *Science Advances*, 7(7):eabe7275, February 2021. Publisher: American Association for the Advancement of Science.

[10] R. Barer and S. Tkaczyk. Refractive Index of Concentrated Protein Solutions. *Nature*, 173(4409):821–822, May 1954.

[11] R. Barer and S. Joseph. Refractometry of Living Cells: Part I. Basic Principles. *Journal of Cell Science*, s3-95(32):399–423, December 1954.

[12] Gabriel Popescu, YoungKeun Park, Niyom Lue, Catherine Best-Popescu, Lauren Deflores, Ramachandra R. Dasari, Michael S. Feld, and Kamran Badizadegan. Optical imaging of cell mass and growth dynamics. *American Journal of Physiology-Cell Physiology*, 295(2):C538–C544, August 2008. Publisher: American Physiological Society.

[13] Abin Biswas, Kyoohyun Kim, Gheorghe Cojoc, Jochen Guck, and Simone Reber. The Xenopus spindle is as dense as the surrounding cytoplasm. *Developmental Cell*, 56(7):967–975.e5, April 2021.

[14] Michael Rubinstein and Ralph H. Colby. *Polymer Physics*. Oxford University Press, New York, NY, 1st edition, June 2003.

### 7 Protein sequences used

**Color key**:
Main protein sequence
TEV protease recognition sequences (cleave between Q | S)
Precission protease recognition sequences (cleave between Q | GP)
Linkers
6xHis-tag
mEGFP-tag
SNAP-tag

>> PGL-3-6xHis-mEGFP

MEANKRQIVEVDGIKSYFFPHLAHYLASNDELLVNNIAQANKLAAFVLGATDKRPSN
EEIAEMILPNDSSAYVLAAGMDVCLILGDDFRPKFDSGAEKLSQLGQAHDLAPIIDDE
KKISMLARKTKLKKSNDAKILQVLLKVLGAEEAEEKFVELSELSSALDLDFDVYVLAK
LLGFASEELQEEIEIIRDNVTDAFEACKPLLKKLMIEGPKIDSVDPFTQLLLTPQEESIE
KAVSHIVARFEEASAVEDDESLVLKSQLGYQLIFLVVRSLADGKRDASRTIQSLMPS
SVRAEVFPGLQRSVFKSAVFLASHIIQVFLGSMKSFEDWAFVGLAEDLESTWRRRA
IAELLKKFRISVLEQCFSQPIPLLPQSELNNETVIENVNNALQFALWITEFYGSESEKK
SLNQLQFLSPKSKNLLVDSFKKFAQGLDSKDHVNRIIESLEKSSSSEPSATAKQTTT
SNGPTTVSTAAQVVTVEKMPFSRQTIPCEGTDLANVLNSAKIIGESVTVAAHDVIPE
KLNAEKNDNTPSTASPVQFSSDGWDSPTKSVALPPKISTLEEEQEEDTTITKVSPQP
QERTGTAWGSGDATPVPLATPVNEYKVSGFGAAPVASGFGQFASSNGTSGRGSY
GGGRGGDRGGRGAYGGDRGRGGSGDGSRGYRGGDRGGRGSYGEGSRGYQG
GRAGFFGGSRGGSSRENLYFQSSAHHHHHHVMVSKGEELFTGVVPILVELDGDV
NGHKFSVSGEGEGDATYGKLTLKFICTTGKLPVPWPTLVTTLTYGVQCFSRYPDHM
KQHDFFKSAMPEGYVQERTIFFKDDGNYKTRAEVKFEGDTLVNRIELKGIDFKEDG
NILGHKLEYNYNSHNVYIMADKQKNGIKVNFKIRHNIEDGSVQLADHYQQNTPGDG
PVLLPDNHYLSTQSALSKDPNEKRDHMVLKEFVTAAGITLGMDELYKGA

>> PGL-3 untagged

MEANKRQIVEVDGIKSYFFPHLAHYLASNDELLVNNIAQANKLAAFVLGATDKRPSN
EEIAEMILPNDSSAYVLAAGMDVCLILGDDFRPKFDSGAEKLSQLGQAHDLAPIIDDE
KKISMLARKTKLKKSNDAKILQVLLKVLGAEEAEEKFVELSELSSALDLDFDVYVLAK
LLGFASEELQEEIEIIRDNVTDAFEACKPLLKKLMIEGPKIDSVDPFTQLLLTPQEESIE
KAVSHIVARFEEASAVEDDESLVLKSQLGYQLIFLVVRSLADGKRDASRTIQSLMPS
SVRAEVFPGLQRSVFKSAVFLASHIIQVFLGSMKSFEDWAFVGLAEDLESTWRRRA
IAELLKKFRISVLEQCFSQPIPLLPQSELNNETVIENVNNALQFALWITEFYGSESEKK
SLNQLQFLSPKSKNLLVDSFKKFAQGLDSKDHVNRIIESLEKSSSSEPSATAKQTTT
SNGPTTVSTAAQVVTVEKMPFSRQTIPCEGTDLANVLNSAKIIGESVTVAAHDVIPE
KLNAEKNDNTPSTASPVQFSSDGWDSPTKSVALPPKISTLEEEQEEDTTITKVSPQP
QERTGTAWGSGDATPVPLATPVNEYKVSGFGAAPVASGFGQFASSNGTSGRGSY
GGGRGGDRGGRGAYGGDRGRGGSGDGSRGYRGGDRGGRGSYGEGSRGYQG
GRAGFFGGSRGGSSRENLYFQ

>> SNAP-TAF15(RBD) (residues 181-589)

GPMDKDCEMKRTTLDSPLGKLELSGCEQGLHRIIFLGKGTSAADAVEVPAPAAVLG
GPEPLMQATAWLNAYFHQPEAIEEFPVPALHHPVFQQESFTRQVLWKLLKVVKFG
EVISYSHLAALAGNPAATAAVKTALSGNPVPILIPCHRVVQGDLDVGGYEGGLAVKE
WLLAHEGHRLGKPGLGGSSSGRENLYFQGAAAGRGRGGYDKDGRGPMTGSSGG
DRGGFKNFGGHRDYGPRTDADSESDNSDNNTIFVQGLGEGVSTDQVGEFFKQIGII
KTNKKTGKPMINLYTDKDTGKPKGEATVSFDDPPSAKAAIDWFDGKEFHGNIIKVSF
ATRRPEFMRGGGSGGGRRGRGGYRGRGGFQGRGGDPKSGDWVCPNPSCGNM
NFARRNSCNQCNEPRPEDSRPSGGDFRGRGYGGERGYRGRGGRGGDRGGYGG
DRSGGGYGGDRSSGGGYSGDRSGGGYGGDRSGGGYGGDRGGGYGGDRGGG
YGGDRGGGYGGDRGGYGGDRGGGYGGDRGGYGGDRGGYGGDRGGYGGDRG
GYGGDRSRGGYGGDRGGGSGYGGDRSGGYGGDRSGGGYGGDRGGGYGGDR
GGYGGKMGGRNDYRNDQRNRPYGAP

>> mEGFP-TAF15(RBD) (residues 181-589)

GPMVSKGEELFTGVVPILVELDGDVNGHKFSVSGEGEGDATYGKLTLKFICTTGKL
PVPWPTLVTTLTYGVQCFSRYPDHMKQHDFFKSAMPEGYVQERTIFFKDDGNYKT
RAEVKFEGDTLVNRIELKGIDFKEDGNILGHKLEYNYNSHNVYIMADKQKNGIKVNF
KIRHNIEDGSVQLADHYQQNTPIGDGPVLLPDNHYLSTQSKLSKDPNEKRDHMVLL
EFVTAAGITLGMDELYKGSSSGRENLYFQGAAAGRGRGGYDKDGRGPMTGSSGG
DRGGFKNFGGHRDYGPRTDADSESDNSDNNTIFVQGLGEGVSTDQVGEFFKQIGII
KTNKKTGKPMINLYTDKDTGKPKGEATVSFDDPPSAKAAIDWFDGKEFHGNIIKVSF
ATRRPEFMRGGGSGGGRRGRGGYRGRGGFQGRGGDPKSGDWVCPNPSCGNM
NFARRNSCNQCNEPRPEDSRPSGGDFRGRGYGGERGYRGRGGRGGDRGGYGG
DRSGGGYGGDRSSGGGYSGDRSGGGYGGDRSGGGYGGDRGGGYGGDRGGG
YGGDRGGGYGGDRGGYGGDRGGGYGGDRGGYGGDRGGYGGDRGGYGGDRG
GYGGDRSRGGYGGDRGGGSGYGGDRSGGYGGDRSGGGYGGDRGGGYGGDR
GGYGGKMGGRNDYRNDQRNRPYGAP

>> FUS-mEGFP

GPAAAMASNDYTQQATQSYGAYPTQPGQGYSQQSSQPYGQQSYSGYSQSTDTS
GYGQSSYSSYGQSQNTGYGTQSTPQGYGSTGGYGSSQSSQSSYGQQSSYPGY
GQQPAPSSTSGSYGSSSQSSSYGQPQSGSYSQQPSYGGQQQSYGQQQSYNPP
QGYGQQNQYNSSSGGGGGGGGGGNYGQDQSSMSSGGGSGGGYGNQDQSGG
GGSGGYGQQASDRGGRGRGGSGGGGGGGGGYNRSSGGYEPRGRGGGRGG
RGGMGGSDRGGFNKFGGPRDQGSRHDSEQDNSDNNTIFVQGLGENVTIESVADY
FKQIGIIKTNKKTGQPMINLYTDRETGKLKGEATVSFDDPPSAKAAIDWFDGKEFSG
NPIKVSFATRRADFNRGGGNGRGGRGRGGPMGRGGYGGGGSGGGGRGGFPSG
GGGGGGQQRAGDWKCPNPTCENMNFSWRNECNQCKAPKPDGPGGGPGGSHM
GGNYGDDRRGGRGGYDRGGYRGRGGDRGGFRGGRGGGDRGGFGPGKMDSR
GEHRQDRRERPYGAPGSAGSAAGSGMVSKGEELFTGVVPILVELDGDVNGHKFS
VSGEGEGDATYGKLTLKFICTTGKLPVPWPTLVTTLTYGVQCFSRYPDHMKQHDFF
KSAMPEGYVQERTIFFKDDGNYKTRAEVKFEGDTLVNRIELKGIDFKEDGNILGHKL
EYNYNSHNVYIMADKQKNGIKVNFKIRHNIEDGSVQLADHYQQNTPIGDGPVLLPD
NHYLSTQSKLSKDPNEKRDHMVLLEFVTAAGITLGMDELYKLEVLFQ

>> BSA (Uniprot ID P02769, FT CHAIN residues 25-607)
DTHKSEIAHRFKDLGEEHFKGLVLIAFSQYLQQCPFDEHVKLVNELTEFAKTCVADE
SHAGCEKSLHTLFGDELCKVASLRETYGDMADCCEKQEPERNECFLSHKDDSPDL
PKLKPDPNTLCDEFKADEKKFWGKYLYEIARRHPYFYAPELLYYANKYNGVFQECC
QAEDKGACLLPKIETMREKVLASSARQRLRCASIQKFGERALKAWSVARLSQKFPK
AEFVEVTKLVTDLTKVHKECCHGDLLECADDRADLAKYICDNQDTISSKLKECCDKP
LLEKSHCIAEVEKDAIPENLPPLTADFAEDKDVCKNYQEAKDAFLGSFLYEYSRRHP
EYAVSVLLRLAKEYEATLEECCAKDDPHACYSTVFDKLKHLVDEPQNLIKQNCDQF
EKLGEYGFQNALIVRYTRKVPQVSTPTLVEVSRSLGKVGTRCCTKPESERMPCTED
YLSLILNRLCVLHEKTPVSEKVTKCCTESLVNRRPCFSALTPDETYVPKAFDEKLFTF
HADICTLPDTEKQIKKQTALVELLKHKPKATEEQLKTVMENFVAFVDKCCAADDKEA
CFAVEGPKLVVSTQTALA
